# Violating the normality assumption may be the lesser of two evils

**DOI:** 10.1101/498931

**Authors:** Ulrich Knief, Wolfgang Forstmeier

## Abstract

When data are not normally distributed (e.g. skewed, zero-inflated, binomial, or count data) researchers are often uncertain whether it may be legitimate to use tests that assume Gaussian errors (e.g. regression, *t*-test, ANOVA, Gaussian mixed models), or whether one has to either model a more specific error structure or use randomization techniques.

Here we use Monte Carlo simulations to explore the pros and cons of fitting Gaussian models to non-normal data in terms of risk of type I error, power and utility for parameter estimation.

We find that Gaussian models are remarkably robust to non-normality over a wide range of conditions, meaning that *P*-values remain fairly reliable except for data with influential outliers judged at strict alpha levels. Gaussian models also perform well in terms of power and they can be useful for parameter estimation but usually not for extrapolation. Transformation of data before analysis is often advisable and visual inspection for outliers and heteroscedasticity is important for assessment. In strong contrast, some non-Gaussian models and randomization techniques bear a range of risks that are often insufficiently known. High rates of false-positive conclusions can arise for instance when overdispersion in count data is not controlled appropriately or when randomization procedures ignore existing non-independencies in the data.

Overall, we argue that violating the normality assumption bears risks that are limited and manageable, while several more sophisticated approaches are relatively error prone and difficult to check during peer review. Hence, as long as scientists and reviewers are not fully aware of the risks, science might benefit from preferentially trusting Gaussian mixed models in which random effects account for non-independencies in the data in a transparent way.

**Tweetable abstract:** Gaussian models are remarkably robust to even dramatic violations of the normality assumption.

## Introduction

In the biological, medical and social sciences, the validity or importance of research findings is generally assessed via statistical significance tests. Significance tests ensure the trustworthiness of scientific results and should reduce the amount of random noise entering the scientific literature. Brunner and Austin (2009) even regard this as the “primary function of statistical hypothesis testing in the discourse of science”. However, the validity of parametric significance tests may depend on whether model assumptions are violated (Gelman & Hill 2007; Zuur *et al.* 2009). In a growing body of literature, researchers express their concerns about irreproducible results (Open Science Collaboration 2015; Ebersole *et al.* 2016; Camerer *et al.* 2018; Silberzahn *et al.* 2018) and it has been argued that the inappropriate use of statistics is a leading cause of irreproducible results (Forstmeier, Wagenmakers & Parker 2017). Yet researchers may often be uncertain about which statistical practices enable them to answer their scientific questions effectively and which might be regarded as error prone.

One of the most widely known assumptions of parametric statistics is the assumption that errors (model residuals) are normally distributed (Lumley *et al.* 2002). This “normality assumption” underlies the most commonly used tests for statistical significance, that is linear models “lm” and linear mixed models “lmm” with Gaussian error, which includes the often more widely known techniques of regression, *t*-test and ANOVA. However, empirical data often deviates considerably from normality, and may even be categorical such as binomial or count data. Recent advances in statistical modelling appear to have solved this problem, because it is now possible to fit generalized linear mixed models “glmm” with a variety of error distributions (e.g. binomial, Poisson, zero-inflated Poisson, negative binomial; O’Hara 2009; Harrison *et al.* 2018) or to use a range of randomization techniques such as bootstrapping (Good 2005) in order to obtain *P*-values and confidence intervals for parameter estimates from data that does not comply with any of those distributions.

While these developments have supplied experts in statistical modelling with a rich and flexible toolbox, we here argue that these new tools also have created substantial damage, because they come with a range of pitfalls that are often not sufficiently understood by a large majority of scientists who are not outspoken experts in statistics, but who nevertheless implement the tools. The diversity of possible mistakes is so large and sometimes specific to certain software applications that we only want to provide some examples that we have repeatedly come across (see Box 1). Our examples include failure to account for overdispersion in glmms with Poisson errors (Harrison 2014; Ives 2015; Forstmeier, Wagenmakers & Parker 2017), inadequate resampling in bootstrapping techniques (e.g. Ihle *et al.* 2019; Santema, Schlicht & Kempenaers 2019), as well as problems with pseudoreplication due to issues with model convergence (Barr *et al.* 2013; Forstmeier, Wagenmakers & Parker 2017; Arnqvist 2020). These issues may lead to anticonservative *P*-values and hence a high risk of false positive claims.

In light of these difficulties we here want to argue whether it may often be “the lesser of two evils” when researchers fit conventional Gaussian (mixed) models to non-normal data, because, as we will show, Gaussian models are remarkably robust to non-normality, ensuring that type I errors (false-positive conclusion) are kept at the desired low rate. Hence, we argue that for the key purpose of limiting type I errors it may often be fully legitimate to model binomial or count data in Gaussian models, and we also would like to raise awareness of some of the pitfalls inherent to non-Gaussian models.

### Box 1

Examples of specialized techniques that may result in biased parameter estimates or in a high rate of false-positive findings due to unrecognized problems of pseudoreplication

A. Many researchers, being concerned about fitting an “inappropriate” Gaussian model, hold the believe that binomial data always requires modelling a binomial error structure, and that count data mandates modelling a Poisson-like process. Yet, what they consider to be “more appropriate for the data at hand” may often fail to acknowledge the non-independence of events in count data (Harrison 2014; Harrison 2015; Ives 2015; Forstmeier, Wagenmakers & Parker 2017). For instance, in a study of butterflies choosing between two species of host plants for egg laying, an individual butterfly may first sit down on species A and deposit a clutch of 50 eggs, followed by a second landing on species B where another 50 eggs are laid. If we characterize the host preference for species A of this individual by the total number of eggs deposited (*p*(A) = 0.5, *N* = 100) we obtain a highly anticonservative estimate of uncertainty (95% CI for *p*(A): 0.398–0.602), while if we base our preference estimate on the number of landings (*p*(A) = 0.5, *N* = 2) we obtain a much more appropriate confidence interval (95% CI for *p*(A): 0.013–0.987). Even some methodological “how-to” guides (e.g. Fordyce *et al.* 2011; Ramsey & Schafer 2013; Harrison *et al.* 2018) forgot to clearly explain that it is absolutely essential to model the non-independence of events via random effects or overdispersion parameters (Zuur *et al.* 2009; Harrison 2014; Harrison 2015; Ives 2015). Unfortunately, non-Gaussian models with multiple random effects often fail to reach model convergence (e.g. Brooks *et al.* 2017), which often lets researchers settle for a model that ignores non-independence and yields estimates with inappropriately high confidence and statistical significance (Barr *et al.* 2013; Forstmeier, Wagenmakers & Parker 2017; Arnqvist 2020).
B. When observational data do not comply with any distributional assumption, randomization techniques like bootstrapping seem to offer an ideal solution for working out the rate at which a certain estimate arises by chance alone (Good 2005). However, such resampling can also be risky in terms of producing false-positive findings if the data is structured (temporal autocorrelation, random effects; e.g. Ihle *et al.* 2019) and if this structure is not accounted for in the resampling regime (blockwise bootstrap; e.g. Önöz & Bayazit 2012). Specifically, there is the risk that non-independence introduces a strong pattern in the observed data, but, in the simulated data, comparably strong patterns do not emerge because the confounding non-independencies were broken up (Ihle *et al.* 2019). We argue that pseudoreplication is a well-known problem that has been solved reasonably well within the framework of mixed models, and the consideration or neglect of essential random effects can be readily judged from tables that present the model output. In contrast, the issue of pseudoreplication is more easily overlooked in studies that implement randomization tests, where the credibility of findings hinges on details of the resampling procedure that are not understood by the majority of readers.
C. When distributions of counts contain a high fraction of zeroes, many researchers 105 think that this issue can be fixed by specifying a zero-inflated model with Poisson or negative binomial error structure. However, they may not be aware of the concept that underlies such models and hence may not understand that such a model, depending on the distribution of the non-zero values, may effectively treat all zeroes as missing values rather than as valid data. This could yield biased parameter estimates in those cases where zeroes represent a valid phenotype rather than a case of missing information. In contrast, such zero-inflated models are ideal when trying to separate two processes, one that is responsible for the occurrence of (some of the) zeroes and one that is responsible for variation in counts (possibly including some zeroes; Brooks *et al*. 2017).

### A wide range of opinions about violating the normality assumption

Throughout the scientific literature, linear models are typically said to be robust to the violation of the normality assumption when it comes to hypothesis testing and parameter estimation as long as outliers are handled properly (Box & Watson 1962; Miller 1986; Ali & Sharma 1996; Lumley *et al.* 2002; Gelman & Hill 2007; Zuur, Ieno & Elphick 2010; Ramsey & Schafer 2013; Williams, Grajales & Kurkiewicz 2013; Puth, Neuhauser & Ruxton 2014; Warton *et al.* 2016), yet authors seem to differ notably in their opinion on how serious we should take the issue of non-normality.

At one end of the spectrum, Gelman and Hill (2007) write “The regression assumption that is generally *least* important is that the errors are normally distributed” and “Thus, in contrast to many regression textbooks, we do not recommend diagnostics of the normality of regression residuals” (p. 46). At the other end of the spectrum, Osborne and Waters (2002) highlight four assumptions of regression that researchers should *always* test, the first of which is the normality assumption. They write “Non-normally distributed variables (highly skewed or kurtotic variables, or variables with substantial outliers) can distort relationships and significance tests”. And since only few research articles report having tested the assumptions underlying the tests presented, Osborne and Waters (2002) worry that they are “forced to call into question the validity of many of these results, conclusions and assertions”.

Between those two ends of the spectrum, many authors adopt a cautious attitude, and regard models that violate the distributional assumptions as ranging from “risky” to “not appropriate”, hence pleading for the use of transformations (e.g. Miller 1986; Bishara & Hittner 2012; Puth, Neuhauser & Ruxton 2014), non-parametric statistics (e.g. Miller 1986), randomization procedures (e.g. Bishara & Hittner 2012; Puth, Neuhauser & Ruxton 2014), or generalized linear models where the Gaussian error structure can be changed to other error structures (e.g. Poisson, binomial, negative binomial) that may better suit the nature of the data at hand (O’Hara 2009; O’Hara & Kotze 2010; Fordyce *et al.* 2011; Warton & Hui 2011; Szöcs & Schäfer 2015; Warton *et al.* 2016; Harrison *et al.* 2018). The latter suggestion, however, may bear a much more serious risk: while Gaussian models are generally accepted to be fairly robust to non-normal errors (here and in the following, we mean by “robust” ensuring a reasonably low rate of type I errors), Poisson models are highly sensitive if their distributional assumptions are violated (see Box 1), leading to a substantially increased risk of type I errors if overdispersion remains unaccounted for (Warton & Hui 2011; Ives 2015; Szöcs & Schäfer 2015; Warton *et al.* 2016).

In face of this diverse literature, it is rather understandable that empirical researchers are largely uncertain about the importance of adhering to the normality assumption in general, and about how much deviation and which form of deviation might be tolerable under which circumstances (in terms of sample size and significance level threshold). With the present article we hope to provide clarification and guidance.

We here use Monte Carlo simulations to explore how violations of the normality assumption affect the probability of drawing false-positive conclusions (the rate of type I errors), because these are the greatest concern in the current reliability crisis (Open Science Collaboration 2015). We aim at deriving simple rules of thumb, which researchers can use to judge whether the violation may be tolerable and whether the *P*-value can be trusted. We also assess the effects of violating the normality assumption in terms of bias and precision on parameter estimation. Furthermore, we provide an R package (“TrustGauss”) that researchers can use to explore the effect of specific distributions on the reliability of *P*-values and parameter estimates.

Counter to intuition, but consistent with a considerable body of literature (Box & Watson 1962; Miller 1986; Ali & Sharma 1996; Lumley *et al.* 2002; Gelman & Hill 2007; Zuur, Ieno & Elphick 2010; Ramsey & Schafer 2013; Williams, Grajales & Kurkiewicz 2013; Puth, Neuhauser & Ruxton 2014; Warton *et al.* 2016) we find that violations of the normality of residuals assumption are rarely problematic for hypothesis testing and parameter estimation, and we argue that the commonly recommended solutions may bear greater risks than the one to be solved.

### The linear regression model and its assumptions

At this point we need to briefly introduce the notation for the model of least squares linear regression. In its simplest form, it can be formulated as *Y*_*i*_ = *a* + *b* × *X*_*i*_ + *e*_*i*_, where each element of the dependent variable *Y*_*i*_ is linearly related to the predictor *X*_*i*_ through the regression coefficient *b* (slope) and the intercept *a*. *e*_*i*_ is the error or residual term, which describes the deviations (residuals) of the actual from the true unobserved (error) or the predicted (residual) *Y*_*i*_ and whose sum equals zero (Sokal & Rohlf 1995; Gelman & Hill 2007). An *F*-test is usually employed for testing the significance of regression models (Ali & Sharma 1996).

Basic statistics texts introduce (about) five assumptions that need to be met for interpreting all estimates from linear regression models safely (**Box 2**: validity, independence, linearity, homoscedasticity of the errors and normality of the errors; Gelman & Hill 2007). Out of these assumptions, normally distributed errors are generally assumed to be the least important (yet probably the most widely known; Lumley *et al.* 2002; Gelman & Hill 2007). Deviations from normality usually do not bias regression coefficients (Ramsey & Schafer 2013; Williams, Grajales & Kurkiewicz 2013) or impair hypothesis testing (no inflated type I error rate, e.g. Bishara & Hittner 2012; Ramsey & Schafer 2013; Puth, Neuhauser & Ruxton 2014; Ives 2015; Szöcs & Schäfer 2015; Warton *et al.* 2016) even at relatively small sample sizes. With large sample sizes ≥ 500 the Central Limit Theorem guarantees that the regression coefficients are on average normally distributed (Ali & Sharma 1996; Lumley *et al.* 2002).

#### Box 2.

Five assumptions of regression models: validity, independence, linearity, homoscedasticity of the errors and normality of the errors (Gelman & Hill 2007). Three of these criteria are concerned with the dependent variable *Y*, or — to be more precise — the regression error *e* (assumptions 2, 4 and 5, see below). The predictor *X* is often not considered, although *e* is supposed to be normal and of equal magnitude at every value of *X*.

1. *Validity* is not a mathematical assumption *per se* but it still poses “the most challenging step in the analysis” (Gelman & Hill 2007), namely that regression should enable the researcher to answer the scientific question at hand (Kass *et al.* 2016).
2. Each value of the dependent variable *Y* is influenced by only a single value of the predictor *X*, meaning that all observations and regression errors *e*_*i*_ are *independent* (Quinn & Keough 2002). Dependence among observations commonly arises either through cluster (i.e. data collected on subgroups) or serial effects (i.e. data collected in temporal or spatial proximity; Ramsey & Schafer 2013). We will discuss the independence assumption later because it is arguably the riskiest to violate in terms of producing type I errors (Zuur *et al.* 2009; see “A word of caution”).
3. The dependent variable *Y* and the predictors should be *linearly* (and additively) related through the regression coefficient *b*. That being said, quadratic or higher-order polynomial relationships can also be accommodated by squaring or raising the predictor variable *X* to a higher power, because *Y* is still modelled as a linear function through the regression coefficient (Williams, Grajales & Kurkiewicz 2013).
4. The variance in the regression error *e* (or the spread of the response around the regression line) is constant across all values of the predictor *X*, i.e. the samples are *homoscedastic*. Deviations from homoscedasticity will not bias parameter estimates of the regression coefficient *b* (Gelman & Hill 2007). Slight deviations are thought to have only little effects on hypothesis testing (Osborne & Waters 2002) and can often be dealt with by weighted regression, mean-variance stabilizing data transformations (e.g. log-transformation) or estimation of heteroscedasticity-robust standard errors (Huber 1967; White 1980; Miller 1986; Zuur *et al.* 2009; see “A word of caution” for further discussion).
5. The errors of the model should be normally distributed (*normality* assumption), which should be tested via inspecting the distribution of the model residuals *e* (Zuur, Ieno & Elphick 2010). Both visual approaches (probability or QQ-plots) and formal statistical tests (Shapiro-Wilk) are commonly applied. Formal tests for normality have been criticized because they have low power at small sample sizes and almost always yield significant deviations from normality at large sample sizes (Ghasemi & Zahediasl 2012). Thus, researchers are mostly left with their intuition to decide how severely the normality assumption is violated and how robust regression is to such violations. A researcher who examines the effect of a single treatment on multiple dependent variables (e.g. health parameters) may adhere strictly to the normality assumption and thus switch forth and back between reporting parametric and non-parametric test statistics depending on how strongly the trait of interest deviates from normality, rendering a comparison of effect sizes difficult.

Importantly, the robustness of regression methods to deviations from normality of the regression errors *e* does not only depend on sample size, but also on the distribution of the predictor *X* (Box & Watson 1962; Mardia 1971). Specifically, when the predictor variable *X* contains a single outlier, then it is possible that the case coincides with an outlier in *Y*, creating an extreme observation with high leverage on the regression line. This is the only case where statistical significance gets seriously misestimated based on the assumption of Gaussian errors in *Y* which is violated by the outlier in *Y*. This problem has been widely recognized (Box & Watson 1962; Miller 1986; Ali & Sharma 1996; Osborne & Waters 2002; Zuur, Ieno & Elphick 2010; Ramsey & Schafer 2013) leading to the conclusion that Gaussian models are robust as long as there are no outliers that occur in *X* and *Y* simultaneously. Conversely, violations of the normality assumption that do not result in outliers should not lead to elevated rates of type I errors.

Distributions of empirical data may deviate from a Gaussian distribution in multiple ways. Rather than being continuous, data may be discrete, such as integer counts or even binomial character states (yes/no data). Continuous variables may deviate from normality in terms of skewness (showing a long tail on one side), kurtosis (curvature leading to light or heavy tails), and even higher-order moments. All these deviations are generally thought to be of little concern (e.g. Bishara & Hittner 2012), even if they are far from fitting to the bell-shaped curve, such as binomial data (Cochran 1950). However, heavily skewed distributions typically result in outliers, which, depending on the distribution of *X*, can be problematic in terms of type I error rates as just explained above (see also Blair & Lawson 1982). In our simulations we try to representatively cover much of the diversity in possible distributions, in order to provide a broad overview that extends beyond the existing literature. We focus on fairly drastic non-normality because only little bias can be expected from minor violations (Hack 1958; Glass, Peckham & Sanders 1972; Bishara & Hittner 2012; Puth, Neuhauser & Ruxton 2014).

### Simulations to assess effects on *P*-values, power and parameter estimates

To illustrate the consequences of violating the normality assumption, we performed Monte Carlo simulations on five continuous and five discrete distributions that were severely skewed, platy- and leptokurtic or zero-inflated (distributions D0–D9 in Figure 1A left column, Table 1), going beyond previous studies that examined less dramatic violations (Bishara & Hittner 2012; Puth, Neuhauser & Ruxton 2014; Ives 2015; Szöcs & Schäfer 2015; Warton *et al.* 2016) but that are still of biological relevance (Gelman & Hill 2007; Frank 2009; Zuur *et al.* 2009). For example, measures of fluctuating asymmetry are distributed half-normally (distribution D4, Table 1) or survival data can be modelled using a gamma distribution (distribution D9, Table 1). The R-code for generating these distributions can be found in the R package “TrustGauss” in the Supplementary Material, where we also provide the specific parameter settings used for generating distributions D0–D9. Moments of these distributions are provided in Table 1. We explored these 10 distributions across a range of sample sizes (*N* = 10, 25, 50, 100, 250, 500, 1000). Starting with the normal distribution D0 for reference, we sorted the remaining distributions D1–D9 by increasing tendency to produce strong outliers because these are known to be problematic (calculated as the average proportion of data points with Cook’s distance exceeding a critical value (see below) at a sample size of *N* = 10). We used these data both as our dependent variable *Y* and as our predictor variable *X* in linear regression models, yielding 10 × 10 = 100 combinations of *Y* and *X* for each sample size (see **Figure S1** for distributions of the independent variable *Y*, the predictor *X*, and residuals). A detailed documentation of the TrustGauss-functions and their application is provided in the Supplement.

**Table 1.** Description of the 10 simulated distributions of the independent variable *Y* and the predictor *X*.

| Name | Sampling distribution | Mean | Variance | Categories | Degree of zero-inflation | Skewness<sup>†</sup> | Kurtosis<sup>‡</sup> | Arguments in TrustGauss<sup>§</sup> |
| --- | --- | --- | --- | --- | --- | --- | --- | --- |
| D0 | Gaussian | 0 | 1 | - | 0 | $1.9 \times 10^{-5}$ | 3.00 | DistributionY="Gaussian", MeanY.gauss=0, SDY.gauss=1 |
| D1 | Binomial | 0.5 | 0.25 | - | 0 | $6.5 \times 10^{-6}$ | 1.00 | DistributionY="Binomial", zeroLevelY.zero=0.5 |
| D2 | Gaussian with categories and zero-inflation<sup>#</sup> | 0 | 1 | 5 | 0.5 | 0.64 | 2.02 | DistributionY="GaussianZeroCategorical", MeanY.gauss=3, SDY.gauss=1, nCategoriesY.cat=5 |
| D3 | Gaussian with zero-inflation<sup>#</sup> | 0 | 1 | - | 0.5 | 0.45 | 1.69 | DistributionY="GaussianZero", MeanY.gauss=3, SDY.gauss=1, zeroLevelY.zero=0.5 |
| D4 | Absolute Gaussian<sup>#</sup> | 0 | 1 | - | 0 | 1.00 | 3.87 | DistributionY="AbsoluteGaussian", MeanY.gauss=0, SDY.gauss=1 |
| D5 | Student's t | 0 | 2 | - | 0 | 0.01 | 20.71 | DistributionY="StudentsT", DFY.student=4 |
| D6 | Gamma with categories<sup>#</sup> | 10 | 100 | 3 | 0 | 3.45 | 15.09 | DistributionY="GammaCategorical", nCategoriesY.cat=3, ShapeY.gamma=1, ScaleY.gamma=10 |
| D7 | Negative Binomial | 10 | 110 | - | 0 | 2.00 | 9.02 | DistributionY="NegativeBinomial", ShapeY.gamma=1, ScaleY.gamma=10 |
| D8 | Binomial | 0.9 | 0.09 | - | 0 | -2.67 | 8.12 | DistributionY="Binomial", zeroLevelY.zero=0.90 |
| D9 | Gamma | 10 | 1000 | - | 0 | 6.32 | 62.84 | DistributionY="Gamma", ShapeY.gamma=0.1, ScaleY.gamma=100 |

<sup>#</sup> Mean and Variance refer to the distributions prior to adding categories, zero-inflation or taking the absolute values.
<sup>†</sup> Skewness and kurtosis were estimated from the simulated distributions with 50 million data points using the moments R package (v0.14, Komsta & Novomestky 2015).
<sup>§</sup> Here we specified the arguments for the dependent variable $Y$ only. However, the specified values are identical for the independent variable $X$ .

**Figure 1.**
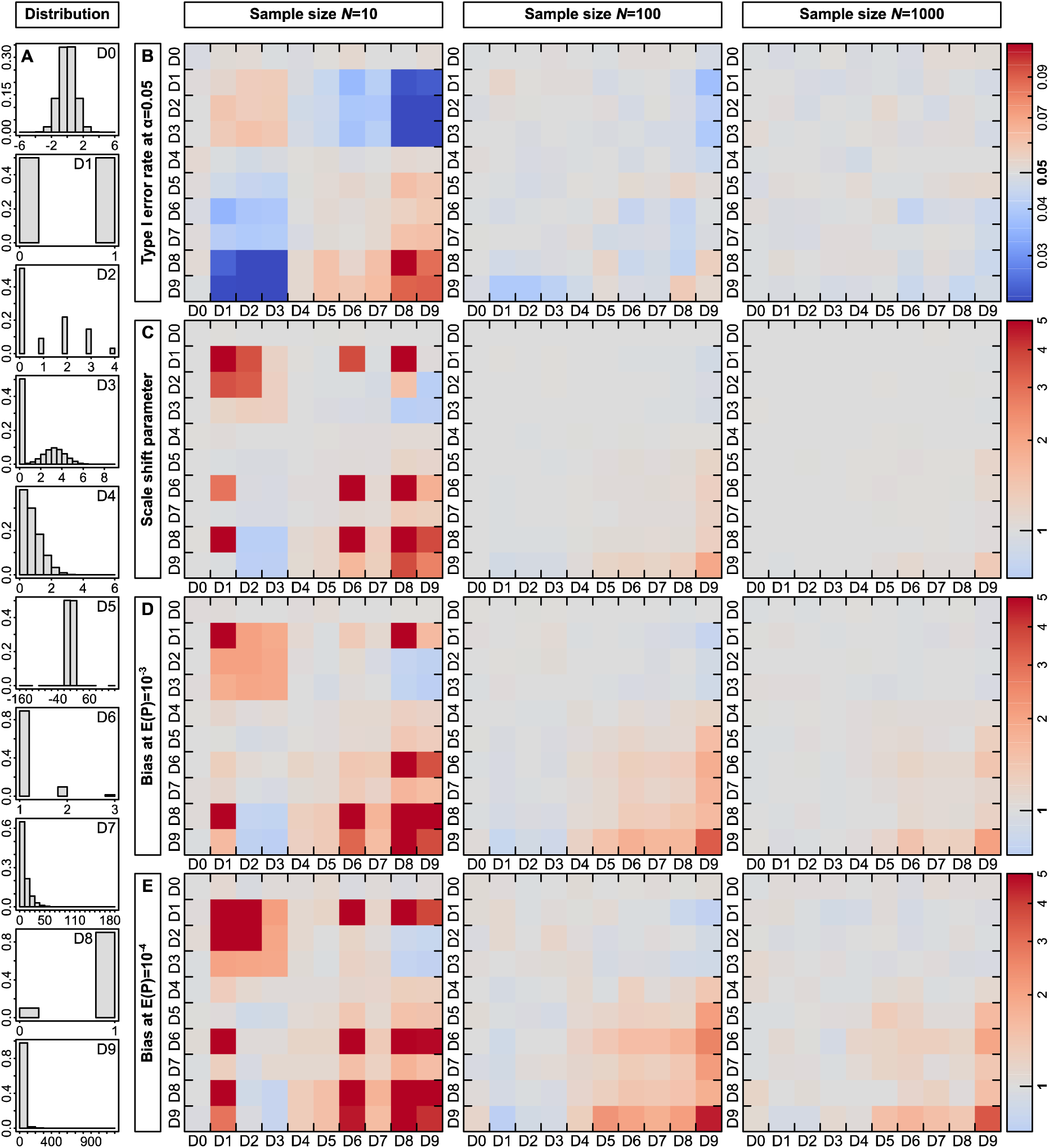
*P*-values from Gaussian linear regression models are in most cases unbiased. (**A**) Overview of the ten different distributions that we simulated. Distributions D0 is Gaussian and all remaining distributions are sorted by their tendency to produce strong outliers. Distributions D1, D2, D6, D7 and D8 are discrete. The numbers D0–D9 refer to the plots in (**B–E**) where on the *Y*-axis the distribution of the dependent variable and on the *X*-axis of the predictor is indicated. (**B**) Type I error rate at an α-level of 0.05 for sample sizes of *N* = 10, 100 and 1000. Red colours represent increased and blue conservative type I error rates. (**C**) Scale shift parameter, (**D**) bias in *P*-values at an expected *P*-value of 10^−3^ and (**E**) bias in *P*-values at an expected *P*-value of 10^−4^.

We assessed the significance of the models via an *F*-test wherever possible and used a likelihood ratio test otherwise. We fitted these models to 50,000 datasets for each combination of the dependent and predictor variable. We did not simulate any effect, which means that both the regression coefficient *b* and the intercept *a* were on average zero. This enabled us to use the frequency of all models that yielded a *P*-value ≤ 0.05 as an estimate of the type I error rate at a significance level (α) of 0.05. The null distribution of *P*-values is uniform on the interval [0,1] and because all *P*-values are independent and identically distributed, we constructed concentration bands using a beta-distribution (cf. Casella & Berger 2002; Knief *et al.* 2017; QQ-plots of expected vs observed *P*-values are depicted in **Figure S1**). We assessed the deviation of observed from expected -log_10_(*P*-values) at an expected exponent value of 3 (*P* = 10^−3^; -log_10_(10^−3^) = 3) and 4 (*P* = 10^−4^) and by estimating the scale shift parameter υ = σ_observed_ / σ_expected_ (Lin 1989), where σ is the standard deviation in -log_10_(*P*-values). We further calculated studentized residuals (*R*), hat values (*H*) and Cook’s distances (*D*) as measures of discrepancy, leverage and influence, respectively, and assessed which proportion exceeded critical values of *R* > 2, *H* > (2 × (*k* + 1)) / *n* and *D* > 4 / (*n* -*k* −1), where *k* is the number of regression slopes and *n* is the number of observations (Zuur, Ieno & Smith 2007).

Since some of the predictor variables were binary rather than continuous, our regression models also comprise the situation of classical two-sample *t*-tests, and we assume that the results would also generalize to the situation of multiple predictor levels (ANOVA), which can be decomposed to multiple binary predictors. To demonstrate that our conclusions from univariate models (involving a single predictor) generalize to the multivariate case (involving several predictors), we fitted the above models with a sample size of *N* = 100 to the same 10 dependent variables with three normally distributed predictors and one additional predictor sampled from the 10 different distributions. We further fitted the above models as mixed-effects models using the lme4 R package (v1.1-14, Bates *et al.* 2015). For that we simulated *N* = 100 independent samples each of which was sampled twice, such that the single random effect “sample ID” explained roughly 30% of the variation in *Y*. We encourage readers to try their own simulations using our R package “TrustGauss”.

We evaluated power, bias and precision of parameter estimates using a sample size of *N* = 10, 100, 1000 and the same 10 distributions of the independent and dependent variables as above. We simulated multivariate data by first Z-transforming the independent variable *Y* and the covariate *X*. We then used an iterative algorithm (SI technique, Ruscio & Kaczetow 2008) that samples from the Z-transformed distributions of *Y* and *X* to introduce a predefined effect size of *r* = 0.15, 0.2 and 0.25 in 50,000 simulations. Additionally, to remove the dominating effect of sample size on power calculations, we calculated the effect size that would be needed to reach a power of 0.5 (rounded to the third decimal) for *N* = 10, 100 and 1000 if *Y* and *X* were normally distributed using the powerMediation R package (v0.2.9, Dupont & Plummer 1998; Qiu 2018). This yielded effect sizes of 0.59, 0.19 and 0.062, respectively. We then introduced effects of such magnitudes with their respective sample sizes in 50,000 simulations. For distribution D6 and the combinations of D8 with D9 we were unable to introduce the predefined effect size also at very large sample sizes (*N* = 100,000) and we removed those from further analyses. We estimated power (β) as the proportion of all simulations that yielded a significant (at α = 0.05 or α = 0.001) regression coefficient *b*. In the case of normally distributed *Y* and *X*, this yielded power estimates that corresponded well with the expectations calculated using the powerMediation R package (v0.2.9, **Table S1**, Dupont & Plummer 1998; Qiu 2018). We used the mean and the coefficient of variation (CV) of the regression coefficient *b* as our measures of bias and precision, respectively. We also assessed interpretability and power of Gaussian versus binomial (mean = 0.75) and Poisson (mean = 1) at a sample size of *N* = 100 by fitting models with a Gaussian, binomial or Poisson error structure in the glms. The effect sizes were chosen such that we reached a power of around 0.5 (see **Table S2** for details on distributions and effect sizes) and models were fitted to 50,000 of such datasets.

## Results

### Effects on P-values

The rate at which linear regression models with Gaussian error structure produced false-positive results (type I errors) was very close to the expected value of 0.05 (Figure 1B). When sample size was high (*N* = 1000), type I error rates ranged only between 0.044 and 0.052, across the 100 combinations of distributions of the dependent variable *Y* and the predictor *X*. Hence, despite of even the most dramatic violations of the normality assumption (see e.g. distributions D8 and D9 in Figure 1A), there was no increased risk of obtaining false-positive results. At *N* = 100, the range was still remarkably narrow (0.037–0.058), and only for very low sample sizes (*N* = 10) we observed 4 out of 100 combinations which yielded notably elevated type I error rates in the range of 0.086 to 0.11. These four cases all involved combinations of the distributions D8 and D9, which yield extreme leverage observations (**Figure S2**). For this low sample size of *N* = 10, there were also cases where type I error rates were clearly too low (down to 0.015, involving distributions D1–D3 where extreme values are rarer than under the normal distribution D0; for details see **Figure S2** and **Table S3**).

Next, we examine the scale shift parameter (Figure 1C) which evaluates the match between observed and expected distributions of *P*-values across the entire range of *P*-values (not only the fraction at the 5% cut-off). Whenever either the dependent variable *Y* or the predictor *X* was normally distributed, the observed and expected *P*-values corresponded very well (first row and first column in Figure 1C). Accordingly, the *P*-values fell within the 95% concentration bands across their entire range (rightmost column in **Figures S1**). This observation was unaffected by sample size (**Table S4**). However, if both the dependent variable *Y* and the predictor *X* were heavily skewed, consistently inflated *P*-values outside the concentration bands occurred, yet this was almost exclusively limited to the case of *N* = 10 (Figure 1C). For larger sample sizes only the most extreme distribution D9 produced somewhat unreliable *P*-values (Figure 1C). This latter effect of unreliable (mostly anti-conservative) *P*-values was most pronounced when judgements were made at a very strict α-level (Figure 1D α = 0.001 and Figure 1E α = 0.0001). At a sample size of *N* = 100, and for α = 0.001, observed -log_10_(*P*-values) were biased maximally 3.36-fold when both *X* and *Y* were sampled from distribution D9. This means that *P*-values of about *P* = 10^−10^ occurred at a rate of 0.001 (*P* = 10^(−3^ × ^3.36)^ = 10^−10.08^; Figure 1D). At *N* = 100, and for α = 0.0001, the bias was maximally 4.54-fold (Figure 1E). Our multivariate and mixed-model simulations confirmed that these patterns are general and also apply to models with multiple predictor variables (**Figure S3**) and to models with a single random intercept (**Figure S4**).

Based on the 100 simulated scenarios that we have constructed, *P*-values from Gaussian models are highly robust to even extreme violation of the normality assumption and can be trusted, except when involving *X* and *Y* distributions with extreme outliers (distribution D9; see also Blair & Lawson 1982). For very small sample sizes, judgements should preferably be made at α = 0.05 (rather than at more strict thresholds) and should also beware of outliers in both *X* and *Y*. The same distributions of the dependent and the independent variable introduced the same type I error rates, meaning that effects were symmetric (Box & Watson 1962). We reference the reader to the “A word of caution” section, where we discuss both the assumption of equal variances of the errors and the effects of non-normality on other applications of linear regression.

### Effects on power and parameter estimates

Power of linear regression models with a Gaussian error structure was only weakly affected by the distributions of *Y* and *X*, whereas sample size and effect size were much more influential (Figure 2B, **Figures S5B**, **S6B**). Power appears to vary notably between distributions when sample size and hence power are small (*N* = 10 in Figure 2B), but this variability rather closely reflects the corresponding type I error rates shown in Figure 1B (Pearson correlation *r* = 0.69 between Figure 1B and **2B** across the *N* = 79 combinations with power estimates at regression coefficient *b* = 0.2 and sample size *N* = 10). To assess the effects of sample size and non-normality on power, we adjusted the regression coefficients such that power stayed constant at 50% for normally distributed *Y* and *X* at sample sizes of *N* = 10, 100 and 1000 (*b* = 0.59, 0.19 and 0.062, respectively, Figure 2C). Then, for *N* = 1000, power was essentially unaffected by the distribution of *Y* and *X*, ranging from 0.48 to 0.52 for all but one combination of *Y* and *X* (β = 0.45 when *Y* and *X* are distributed as D9, that is gamma Γ(0.1, 100), Table 1). In that particular combination, power was not generally reduced but the distribution of *P*-values was shifted, such that power could either be reduced or increased depending on the α-threshold (at α = 0.001 that combination yielded the highest power). At *N* = 100, power varied slightly more (0.44–0.60) but still 87% of all power estimates were between 0.48 and 0.52. Only at a sample size of *N* = 10, power varied considerably between 0.05 and 0.87 (30% of all estimates between 0.48 and 0.52, Figure 2C).

**Figure 2.**
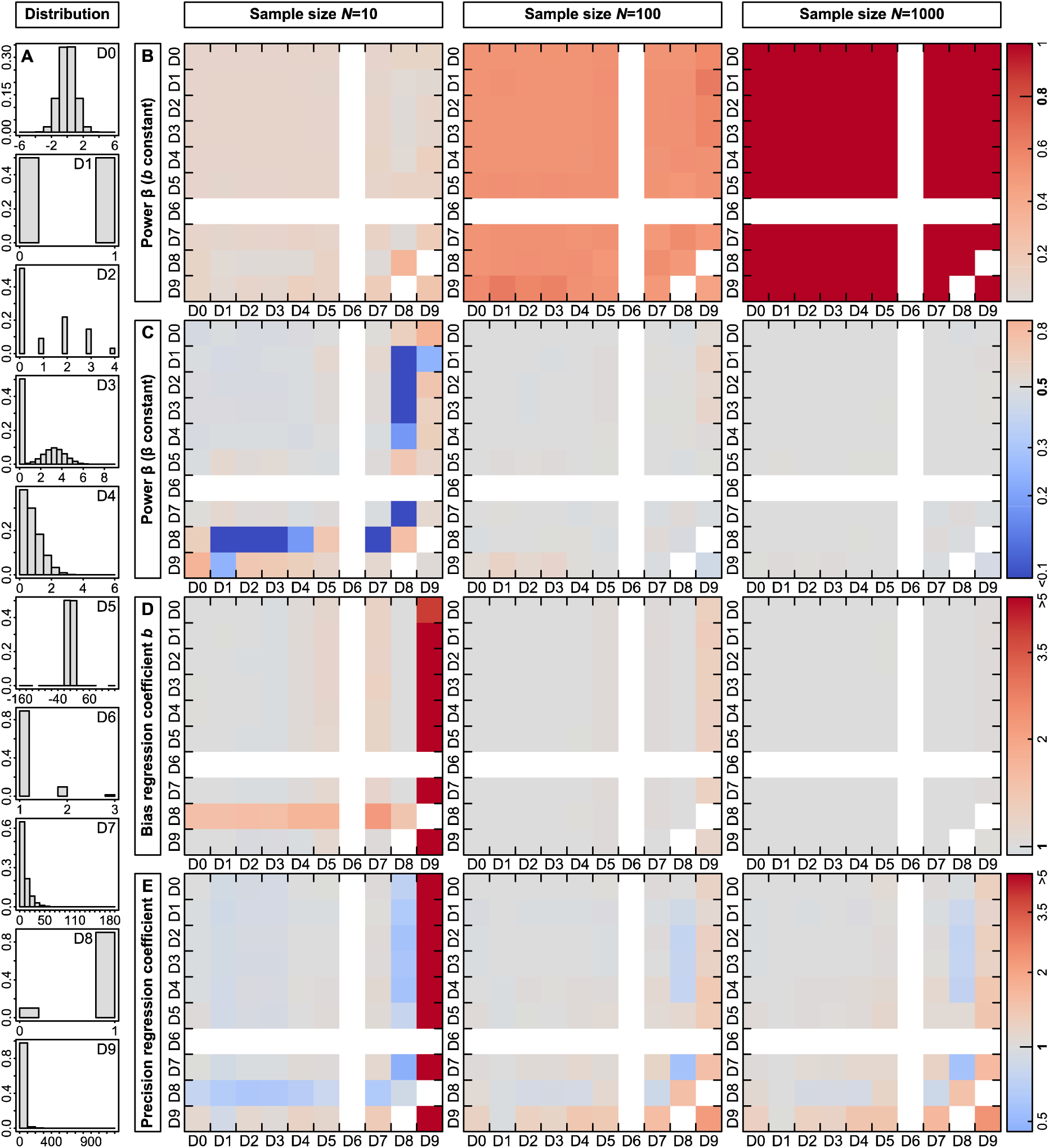
Power, bias and precision of parameter estimates from Gaussian linear regression models are in most cases unaffected by the distributions of the dependent variable *Y* or the predictor *X*. (**A**) Overview of the different distributions that we simulated, which were the same as in **Figure 1**. The numbers D0–D9 refer to the plots in (**B–E**) where on the *Y*-axis the distribution of the dependent variable and on the *X*-axis of the predictor is indicated. (**B**) Power at a regression coefficient *b* = 0.2 for sample sizes of *N* = 10, 100 and 1000. Red colours represent increased power. (**C**) Power at regression coefficients *b* = 0.59, 0.19 and 0.06 for sample sizes of *N* = 10, 100 and 1000, respectively, where the expected power derived from a normally distributed *Y* and *X* is 0.5. Red colours represent increased and blue colours decreased power. (**D**) Bias and (**E**) precision of the regression coefficient estimates at an expected *b* = 0.2 for sample sizes of *N* = 10, 100 and 1000.

For most distributions of *Y* and *X*, regression coefficients were unbiased, which follows from the Lindeberg-Feller Central Limit Theorem (Lumley *et al.* 2002). The strongest bias occurred at a sample size of *N* = 10 and when the distribution of *X* was highly skewed (D9), resulting in such a high frequency of high leverage observations that the Lindeberg-Feller Central Limit Theorem did not hold (**Figure S2**). In the most extreme case, the mean regression coefficients at *N* = 10 were below zero (indicated as additional white squares in **Figure S5D**, **S6D**). However, the bias shrunk to maximally 1.32-fold when the sample size increased to *N* = 100 and to 1.03-fold at a sample size of *N* = 1000 (Figure 2D).

We used the coefficient of variation in regression coefficients as our measure of the precision of parameter estimates. Similar to the pattern in bias, regression coefficients were precise for most distributions of *Y* and *X* and the lowest precision occurred at a sample size of *N* = 10 and when the distribution of *X* was highly skewed (D9). However, there was no gain in precision when increasing the sample size from *N* = 100 to *N* = 1000 (Figure 2E) and precision slightly decreased at larger effect sizes (**Figure S5E**, **S6E**).

We conclude that in our 79 simulated scenarios neither power nor bias or precision of parameter estimates are heavily affected by violations of the normality assumption by both the distributions of the dependent variable *Y* and the predictor *X*, except when involving predictors with extreme outliers (i.e. high leverage, distribution D9). An increase in sample size protects against severely biased parameter estimates but does not make estimates more precise. We provide further advice in the “A word of caution” section.

### Comparison between error distributions

In the previous section, we have shown that Gaussian models are robust to violations of the normality assumption. How do they perform in comparison to Poisson and binomial models and how do Poisson models perform if their distributional assumptions are violated? To address these questions, we fitted glms with a Gaussian, Poisson or binomial error structure to data where the dependent variable *Y* was Gaussian, Poisson or binomial distributed and the predictor variable *X* followed a Gaussian, gamma or binomial distribution. This allowed us to directly compare the effect of the error structure on power, bias and precision of the parameter estimate. Interestingly, models with a Gaussian error structure were largely comparable in terms of power and bias to those fitted using the appropriate error structure. However, parameter estimates were less precise using the Gaussian error structure (Table 2), which argues in favour of the more specialized models for the purpose of parameter estimation.

**Table 2.** Summary of power, bias and precision of parameter estimates and interpretability from 50,000 simulation runs across the six combinations of the dependent variable *Y* and the predictor *X*. Each combination was either fitted using a Gaussian error structure or the appropriate error structure according to the distribution of *Y* (that is either Poisson with a mean of 1 or binomial with a mean of 0.75). The predefined effect was chosen such that a power of around 0.5 was reached (see **Table S2** for details). The column Effect is the mean estimated effect (intercept + slope) after back-transformation.

| Distribution of $Y$ | Distribution of $X$ | Error Distribution | Sample size | Power at $\alpha = 0.05$ | Power at $\alpha = 0.001$ | Mean of slope $b$ | Variance in slope $b$ | CV of slope $b$ | Mean intercept $a$ | Variance in intercept $a$ | CV of intercept $a$ | Effect | Variance in effect |
| --- | --- | --- | --- | --- | --- | --- | --- | --- | --- | --- | --- | --- | --- |
| Poisson | Gaussian | Gaussian | 100 | 0.522 | 0.094 | 0.200 | $9.96 \times 10^{-3}$ | 0.498 | 1.000 | $9.70 \times 10^{-3}$ | 0.098 | 1.201 | 0.023 |
| Poisson | Gaussian | Poisson | 100 | 0.511 | 0.090 | 1.228 | 0.015 | 0.100 | 0.976 | $9.80 \times 10^{-3}$ | 0.101 | 1.195 | 0.022 |
| Binomial | Gaussian | Gaussian | 100 | 0.502 | 0.085 | 0.085 | $1.79 \times 10^{-3}$ | 0.500 | 0.750 | $1.82 \times 10^{-3}$ | 0.057 | 0.835 | $2.84 \times 10^{-3}$ |
| Binomial | Gaussian | Binomial | 100 | 0.504 | 0.091 | 0.617 | $3.63 \times 10^{-3}$ | 0.098 | 0.762 | $2.03 \times 10^{-3}$ | 0.059 | 0.834 | $2.75 \times 10^{-3}$ |
| Poisson | Gamma | Gaussian | 100 | 0.588 | 0.162 | 0.023 | $1.28 \times 10^{-4}$ | 0.502 | 0.776 | $1.28 \times 10^{-4}$ | 0.176 | 0.798 | 0.017 |
| Poisson | Gamma | Poisson | 100 | 0.537 | 0.095 | 1.019 | $7.67 \times 10^{-5}$ | 0.009 | 0.818 | $7.67 \times 10^{-5}$ | 0.142 | 0.833 | 0.013 |
| Binomial | Gamma | Gaussian | 100 | 0.459 | 0.029 | 0.008 | $1.55 \times 10^{-5}$ | 0.481 | 0.669 | $4.12 \times 10^{-3}$ | 0.096 | 0.677 | $3.75 \times 10^{-3}$ |
| Binomial | Gamma | Binomial | 100 | 0.549 | 0.113 | 0.517 | $1.15 \times 10^{-4}$ | 0.021 | 0.634 | $6.87 \times 10^{-3}$ | 0.131 | 0.650 | $5.59 \times 10^{-3}$ |
| Poisson | Binomial | Gaussian | 100 | 0.673 | 0.126 | 0.534 | 0.039 | 0.371 | 0.599 | 0.025 | 0.265 | 1.133 | 0.014 |
| Poisson | Binomial | Poisson | 100 | 0.699 | 0.189 | 1847.624 | $1.70 \times 10^{11}$ | 223.359 | 0.599 | 0.025 | 0.264 | 1.132 | 0.014 |
| Binomial | Binomial | Gaussian | 100 | 0.510 | 0.127 | 0.200 | 0.012 | 0.551 | 0.600 | $9.96 \times 10^{-3}$ | 0.166 | 0.800 | $2.15 \times 10^{-3}$ |
| Binomial | Binomial | Binomial | 100 | 0.491 | 0.094 | 0.717 | 0.011 | 0.146 | 0.600 | 0.010 | 0.167 | 0.800 | $2.16 \times 10^{-3}$ |

More importantly for the reliability of science, and in contrast to Gaussian models, Poisson models are not at all robust to violations of the distribution assumption. For comparison, we fitted the above univariate models involving the five discrete distributions (D1, D2, D6, D7, D8) with a sample size of *N* = 100 using a Poisson error structure (inappropriately). This yielded heavily biased type I error rates (at α = 0.05) in either direction ranging from 0 to as high as 0.55 (Figure 3, right column, **Figures S7**). Yet when also inappropriately modelling these distributions as Gaussian, type I error rates are very close to the nominal level of 0.05 (Figure 3, left column). Controlling for overdispersion in counts through the use of a glmm with an observation-level random effect (Harrison *et al.* 2018) fixed the problem of inflated type I error rates for distributions D2 and D7 (Figure 3, indicated in red) but did not solve the problem of low power for distributions D1, D6, and D8 (Figure 3, indicated in blue). Using a quasi-likelihood method (“Quasipoisson”, Wedderburn 1974) provided unbiased type I error rates, like in the Gaussian models (Figure 3), but this quasi-likelihood method is not available in the mixed-effects package lme4 in R (Bates *et al.* 2015).

**Figure 3.**
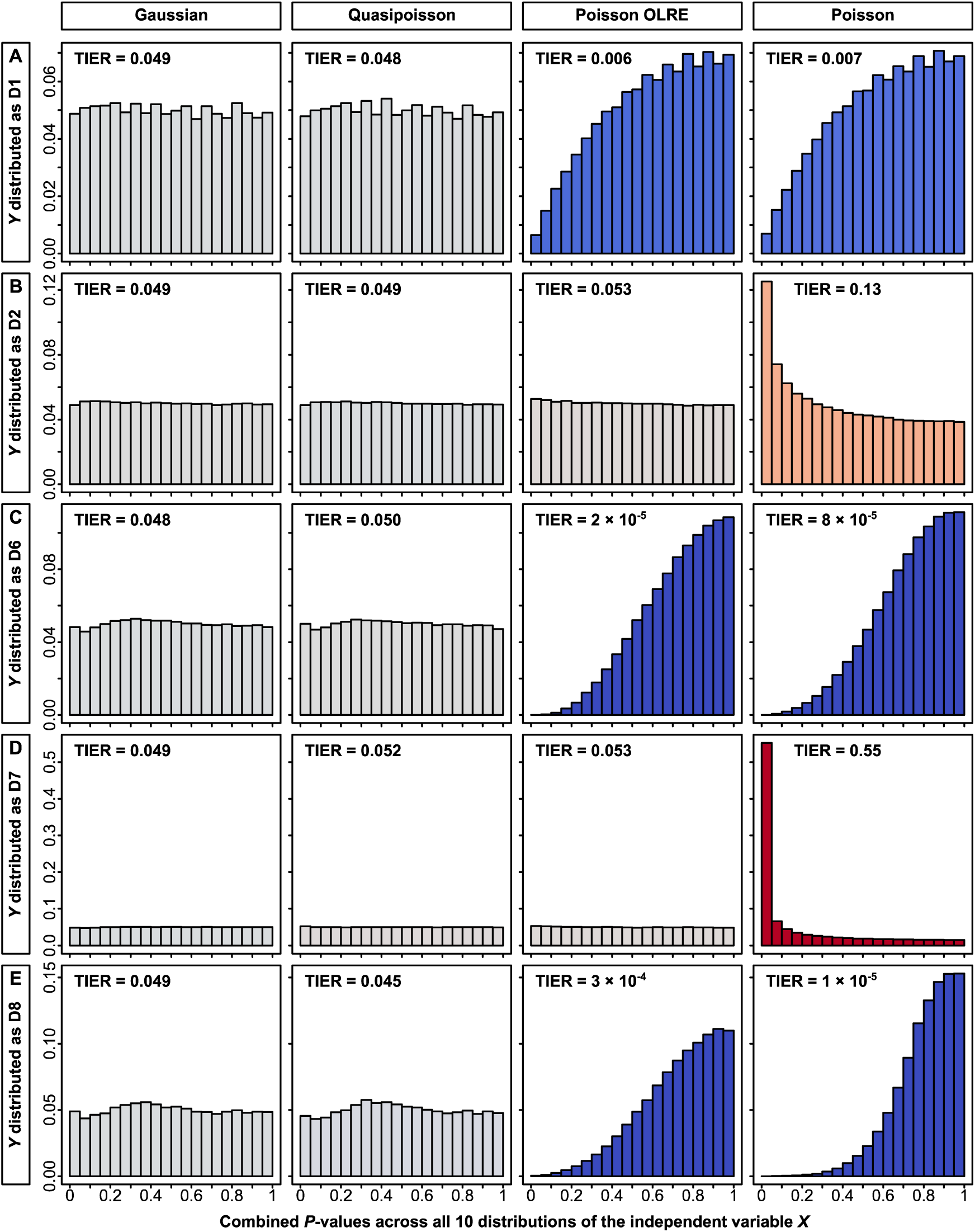
Distribution of observed *P*-values (when the null hypothesis is true) as a function of different model specifications (columns) and different distributions of the dependent variable *Y* (rows **A** to **E**). Each panel was summed up across 10 different distributions of the predictor *X* (500,000 simulations per panel with *N* = 100 data points per simulation). Models were fitted either as glms with a Gaussian error structure that violate the normality assumption (first column), as glms with a Quasipoisson error structure that take overdispersion into account (second column), as glmms with a Poisson error structure and an observation-level random effect (OLRE; Harrison *et al.* 2018) or as glms with a Poisson error structure that violate the assumption of the Poisson distribution. In each panel, TIER indicates the realized type I error rate (across the 10 different predictor distributions), highlighted with a colour scheme as in **Figure 1B** (blue: below the nominal level of 0.05, red: above the nominal level, grey: closely matching the nominal level). The dependent variable *Y* was distributed as (**A**) distribution D1, (**B**) distribution D2, (**C**) distribution D6, (**D**) distribution D7 or (**E**) distribution D8 (see Table 1 and **Figure 1A** for details).

## A word of caution

Our finding that violations of the normality assumption are relatively unproblematic with regard to type I errors should not be misunderstood as a *carte blanche* to violate any assumption of linear models. The probably riskiest assumption to violate (in terms of producing type I errors) is the assumption of independence of data points (Kass *et al.* 2016; Forstmeier, Wagenmakers & Parker 2017), because one tends to overestimate the amount of independent evidence that is provided by the data points, which are not real replicates (hence this is called “pseudoreplication”).

Another assumption that is not to be ignored concerns the homogeneity of variances across the entire range of the predictor variable (Box 1953; Glass, Peckham & Sanders 1972; Miller 1986; McGuinness 2002; Osborne & Waters 2002; Zuur *et al.* 2009; Ramsey & Schafer 2013; Williams, Grajales & Kurkiewicz 2013). Violating this assumption may result in more notable increases of type I errors (compared to what we examined here) at least when the violations are drastic. For instance, when applying a *t*-test that assumes equal variances in both groups to data that come from substantially different variances (e.g. σ_1_^2^/ σ_2_^2^ = 0.1), then high rates of type I errors (e.g. 23%) may be obtained in a situation where sample sizes are unbalanced (*N*_1_ = 15, *N*_2_ = 5), namely when the small sample comes from the more variable group (Glass, Peckham & Sanders 1972; Miller 1986). Also in this example, it is the influence of outliers (small *N* sampled from large variance) that results in misleading *P*-values. We further carried out some extra simulations to explore whether non-normality tends to exacerbate the effects of heteroscedasticity on type I error rates, but we found that normal and non-normal data behaved practically in the same way (see Supplementary Methods and **Table S5**). Hence, heteroscedasticity can be problematic, but this seems to be fairly independent of the distribution of the variables.

Diagnostic plots of model residuals over fitted values can help identifying outliers and recognizing heterogeneity in variances over fitted values. Transformation of variables is often a helpful remedy if one observes that variance strongly increases with the mean. This typically occurs in comparative studies, where e.g. body size of species may span several orders of magnitude (calling for a log-log plot). Most elegantly, heteroscedasticity can be modelled directly, for instance by using the “weights” argument in lme (see Pinheiro & Bates 2000, p. 214), which also enables us to test directly whether allowing for heteroscedasticity increases the fit of the model significantly. Similarly, heteroscedasticity-consistent standard errors could be estimated (Hayes & Cai 2007). For more advice on handling heteroscedasticity see McGuinness (2002).

Another word of caution when running Gaussian models on non-Gaussian data should be expressed when it comes to the interpretation of parameter estimates of models. If the goal of modelling lies in the estimation of parameters (rather than hypothesis testing) then such models should be regarded with caution. First, recall that distributions with extreme outliers are often better characterized by their median than by their mean, which gets pulled away by extreme values. Second, parameter estimates for counts or binomial traits may be acceptable for interpretation when they refer to the average condition (e.g. a typical family having 1.8 children consisting of 50% boys). However, parameter estimates may become nonsensical outside the typical range of data (e.g. negative counts or probabilities). In such cases one might also consider fitting separate models for parameter estimation and for hypothesis testing (Warton *et al.* 2016).

Finally, in the above we were exclusively concerned with associations between variables, that is parameter estimates derived from the whole population of data points. However, sometimes we might be interested in predicting the response of specific individuals in the population and we need to estimate a prediction interval. In that case, a valid prediction interval requires the normality assumption to be fulfilled because it is based directly on the distribution of *Y* (Lumley *et al.* 2002; Ramsey & Schafer 2013).

### The issue of overdispersion in non-Gaussian models

We have shown that Poisson models yielded heavily biased type I error rates (at α = 0.05) in either direction ranging from 0 to as high as 0.55 when their distribution assumption is violated (Figure 3 right column, **Figures S7**). This of course is an inappropriate use of the Poisson model, but still this is not uncommonly found in the scientific literature. Such inflations of type I error rates in glms already have been reported frequently (Young, Campbell & Capuano 1999; Warton & Hui 2011; Ives 2015; Szöcs & Schäfer 2015; Warton *et al.* 2016) and this problem threatens the reliability of research whenever such models are implemented with insufficient statistical expertise.

First, it is absolutely essential to control for overdispersion in the data (that is more extreme counts than expected under a Poisson process), either by using a quasi-likelihood method (“Quasipoisson”) or by fitting an observation level random effect (“OLRE”; Figure 3). Overdispersion may already be present when counts refer to discrete natural entities (for example counts of animals), but may be particularly strong when Poisson errors are less appropriately applied to measurements of areas (e.g. counts of pixels or mm^2^), latencies (e.g. counts of seconds), or concentrations (e.g. counts of molecules). Similarly, there may also be overdispersion in counts of successes versus failures that are being analysed in a binomial model (e.g. fertile versus infertile eggs within a clutch). Failure to account for overdispersion (as in Figure 3B and **3D**) will typically result in very high rates of type I errors (Young, Campbell & Capuano 1999; Warton & Hui 2011; Ives 2015; Szöcs & Schäfer 2015; Warton *et al.* 2016; Forstmeier, Wagenmakers & Parker 2017).

Second, even after accounting for overdispersion, some models may still yield inflated or deflated type I error rates (not observed in our examples of Figure 3), therefore requiring statistical testing via a resampling procedure (Warton & Hui 2011; Ives 2015; Szöcs & Schäfer 2015; Warton *et al.* 2016), but this may also depend on the software used. While several statistical experts have explicitly advocated for such a sophisticated approach to count data (O’Hara 2009; O’Hara & Kotze 2010; Szöcs & Schäfer 2015; Warton *et al.* 2016; Harrison *et al.* 2018), we are concerned about practicability when non-experts have to make decisions about the most adequate resampling procedure, particularly when there are also non-independencies in the data (random effects) that have to be considered. In this field of still developing statistical approaches it seems much easier to get things wrong (and obtain a highly overconfident *P*-value) than to get everything right (Bolker *et al.* 2009).

In summary, we are worried that authors being under pressure to present statistically significant findings will misinterpret type I errors (due to incorrect implementation) optimistically as a true finding and misattribute the gained significance to a presumed gain of power when fitting the “appropriate” error structure (note that such power gains should be quite small; see Table 2 and also Szöcs & Schäfer 2015; Warton *et al.* 2016). Moreover, we worry that sophisticated methods may allow presenting nearly anything as statistically significant (Simmons, Nelson & Simonsohn 2011) because complex methods will only rarely be questioned by reviewers.

### Practical advice

Anti-conservative *P*-values usually do not arise from violating normality in Gaussian models (except for the case of influential outliers), but rather from various kinds of non-independencies in the data (see Box 1). We therefore recommend the Gaussian mixed-effect model as a trustworthy and universal standard tool for hypothesis testing, where transparent reporting of the model’s random effect structure clarifies to the reader which non-independencies in the data were accounted for. Non-normality should not be a strong reason for switching to a more specialized technique, at least not for hypothesis testing, and such techniques should only be used with a good understanding of the risks involved (see Box 1).

To avoid the negative consequences of strong deviations from normality that may occur under some conditions (see Figure 1) it may be most advisable to apply a rank-based inverse normal (RIN) transformation (aka rankit scores, Bliss 1967) to the data, which can approximately normalize most distributional shapes and which effectively minimizes type I errors and maximises statistical power (Bishara & Hittner 2012; Puth, Neuhauser & Ruxton 2014). Note that we have avoided transformations in our study simply to explore the consequences of major non-normality, but we agree with the general wisdom that transformations can mitigate problems with outliers (Osborne & Overbay 2004), heteroscedasticity (McGuinness 2002), and sometimes with interpretability of parameter estimates.

In practice, we recommend the following to referees:

1. When a test assumes Gaussian errors, request a check for influential observations, particularly if very small *P*-values are reported. Consider recommending a RIN-transformation or other transformations for strong deviations from normality.
2. For Poisson models or binomial models of counts, always check whether the issues of overdispersion and resampling are addressed, otherwise request an adequate control for type I errors or verification with Gaussian models.
3. For randomization tests, request clarity about whether observed patterns may be influenced by non-independencies in the data that are broken up by the randomization procedure. If so, ask for possible alternative ways of testing or of randomizing (e.g. blockwise bootstrap).
4. When requesting a switch to more demanding techniques (e.g. non-Gaussian models, randomization techniques), reviewers should accompany this recommendation with sufficient advice, caveats and guidance to ensure a safe and robust implementation.

## Conclusion

If we are interested in statistical hypothesis testing, linear regression models with a Gaussian error structure are generally robust to violations of the normality assumption. Judging *P*-values at the threshold of α = 0.05 is nearly always safe, but if both *Y* and *X* are skewed, we should avoid being overly confident in very small *P*-values and examine whether these result from outliers in both *X* and *Y* (see also Blair & Lawson 1982; Osborne & Overbay 2004). With this caveat in mind, violating the normality assumption is relatively unproblematic and there is much to be gained when researchers follow a standardized way of reporting effect sizes (Lumley *et al.* 2002). This is good news also for those who want to apply models with Gaussian error structure to binomial or count data when models with other structures fail to reach convergence or produce nonsensical estimates (e.g. Ives & Garland 2014; Plaschke *et al.* 2019). While Gaussian models are rarely misleading, other approaches (see examples in Box 1) may bear a non-trivial risk of yielding anti-conservative *P*-values when applied by scientists with limited statistical expertise.

### Data availability

All functions are bundled in an R package named “TrustGauss”. The R package, R scripts, supplementary figures S1, S3, S4 and S7 and the raw simulation outputs are accessible through the Open Science Framework (https://osf.io/r5ym4/?view_only=5d79da4f8b4441e1addf99b0d435a45e).

## Data availability

all scripts bundled in the R package “TrustGauss”

## Acknowledgements

We thank N. Altman, S. Nakagawa, M. Neuhäuser, F. Korner-Nievergelt and H. Schielzeth for helpful discussions and B. Kempenaers and J.B.W. Wolf for their support.

## Author contributions

W.F. and U.K. conceived of the study. U.K. wrote the simulation code. U.K. and W.F. prepared the manuscript.

## Competing interests

The authors declare no competing financial interests.

## Notes

### Competing Interest Statement

The authors have declared no competing interest.

