## Supplementary Material for "Violating the normality assumption may be the lesser of two evils"

#### Index

**Supplementary Methods** 2
**Supplementary Figures** 9
**Supplementary Tables** 16
**References** 23

### Supplementary Methods

**Description of the TrustGauss functions**

The main function of the "TrustGauss" package is TrustGauss() and it can take 29 arguments that are described in the documentation to the function and that can be accessed via

```
> ?TrustGauss
```

TrustGauss() can be used to assess type I error rates of linear regression models that are fitted through a call to the base R function glm(). We here briefly summarize each of the 29 arguments. Default settings for each argument are given in the documentation to the function.

1. `Family`. This argument takes a character input and specifies the error distribution and link function to be used in the generalized linear model. It can take one of the following values: "gaussian", "poisson", "binomial", "quasipoisson", "quasibinomial" or "Gamma". Since the argument is passed directly to the glm() function, the link function can be specified in the standard way, for example as "gaussian(link = 'identity')". See also the glm() function for further details.
2. `nSamples`. This argument takes a numeric integer input, specifying the number of samples/data points to simulate.
3. `nSimulations`. This argument takes a numeric integer input, specifying for how many iterations the simulation will run.
4. `SaveAllOutput`. This argument is Boolean. If it is set to TRUE, all individual data points of the dependent and independent variables are returned in a list. They can be found in "Data" with column names "Dependent", "Cov1", "Cov2", ..., "Fac1", "Fac2", ... (depending on what combination of covariates and factors is specified, see below).
5. `CompareTtest`. This argument is Boolean. If it is set to TRUE, *P*-values are calculated through both the glm() and the t.test() function. This is only valid when a single factor with two levels is fitted as the independent variable with distribution set to "UniformCategorical" (see below).
6. `PlotExample`. This parameter is Boolean. If it is set to TRUE, one example histogram of the distribution of the dependent variable *Y* is plotted.
7. `DistributionY`. This argument takes a character input and specifies the distribution of the dependent variable *Y*. It can take one of the following values: "Gaussian", "GaussianCategorical", "GaussianZero", "AbsoluteGaussian", "Gamma", "GammaCategorical", "GaussianZeroCategorical", "Binomial", "NegativeBinomial", "StudentsT", "Poisson" or "Uniform". In principle, the base R functions for generating randomly distributed Gaussian [rnorm()], Gamma [rgamma()], binomial [rbinom()], negative binomial [rnbinom()], Student's t [rt()], Poisson [rpois()] or uniform [runif()] variables are used. Parameters for all distributions can be specified (see below). "GaussianCategorical" generates normally distributed integers. "GaussianZero" generates a zero-inflated normal distribution. "AbsoluteGaussian" simulates absolute values of a Gaussian distribution. "GaussianZeroCategorical" first generates a zero-inflated normal distribution and then produces categories. "GammaCategorical" generates gamma distributed integers.
8. `DistributionXCov`. This argument takes a character input and specifies the distribution of the independent covariate *X*. It can take the same values as DistributionY. It is also possible to specify multiple different distributions in order to fit more than one covariate (as a vector). Additionally, it can be set to NULL if only factors should be fitted.

9. `DistributionXFac`. This argument takes a character input and specifies the distribution of the independent factor *X*. Only the categorical distributions are valid inputs ("GaussianCategorical", "GammaCategorical", "GaussianZeroCategorical", "Binomial" or "UniformCategorical"). It is also possible to specify multiple different distributions in order to fit more than one factor (as a vector). Additionally, it can be set to NULL if only covariates should be fitted.

The following arguments specify parameters for the distributions of the independent and dependent variables.

10. `MeanY.gauss`. This argument takes a numeric input, specifying the mean of the distribution of *Y*, if DistributionY is set to "Gaussian", "GaussianCategorical", "GaussianZero", "GaussianZeroCategorical" or "AbsoluteGaussian". See also the rnorm() function for further details. The operations of categorization, taking the absolute value or adding zero-inflation are performed after the call to the rnorm() function.
11. `SDY.gauss`. This argument takes a numeric input, specifying the standard deviation of the distribution of *Y*, if DistributionY is set to "Gaussian", "GaussianCategorical", "GaussianZero", "GaussianZeroCategorical" or "AbsoluteGaussian". See also the rnorm() function for further details. The operations of categorization, taking the absolute value or adding zero-inflation are performed after the call to the rnorm() function.
12. `nCategoriesY.cat`. This argument takes a numeric integer input, specifying how many categories are simulated, if DistributionY is set to "GaussianCategorical" or "GammaCategorical".
13. `zeroLevelY.zero`. This argument takes a numeric input, specifying the proportion of data that will be set to 0, if DistributionY is set to "GaussianZero" or "GaussianZeroCategorical".
14. `ShapeY.gamma`. This argument takes a numeric input, specifying the shape parameter k, if DistributionY is set to "Gamma", "GammaCategorical" or "NegativeBinomial". See also the rgamma() and rnbinom() functions for further details. Categorization is performed after the call to the rgamma() function.
15. `ScaleY.gamma`. This argument takes a numeric input, specifying the scale parameter capital theta, if DistributionY is set to "Gamma", "GammaCategorical" or "NegativeBinomial". See also the rgamma() and rnbinom() functions for further details. Categorization is performed after the call to the rgamma() function.
16. `DFY.student`. This argument takes a numeric input, specifying the degrees of freedom, if DistributionY is set to "StudentsT". See also the rt() function for further details.
17. `MinY.uni`. This argument takes a numeric input, specifying the minimum of the distribution, if DistributionY is set to "Uniform". See also the runif() function for further details.
18. `MaxY.uni`. This argument takes a numeric input, specifying the maximum of the distribution, if DistributionY is set to "Uniform". See also the runif() function for further details.
19. `LambdaY.pois`. This argument takes a numeric input, specifying the mean of the distribution, if DistributionY is set to "Poisson". See also the rpois() function for further details.
20. `MeanX.gauss`. This argument takes a numeric input, specifying the mean of the distribution of the independent variable *X*, if DistributionX is set to "Gaussian", "GaussianCategorical", "GaussianZero", "GaussianZeroCategorical" or "AbsoluteGaussian". See also the rnorm() function for further details. See also the rnorm() function for further details. The operations of categorization, taking the absolute value or adding zero-inflation are performed after the call to the rnorm() function.
21. `SDX.gauss`. This argument takes a numeric input, specifying the standard deviation of the distribution of the independent variable *X*, if DistributionX is set to "Gaussian",

"GaussianCategorical", "GaussianZero", "GaussianZeroCategorical" or "AbsoluteGaussian". See also the rnorm() function for further details. See also the rnorm() function for further details. The operations of categorization, taking the absolute value or adding zero-inflation are performed after the call to the rnorm() function.

22. `nCategoriesX.cat`. This argument takes a numeric integer input, specifying how many categories are simulated, if DistributionX is set to "GaussianCategorical" or "GammaCategorical".
23. `zeroLevelX.zero`. This argument takes a numeric input, specifying the proportion of data that will be set to 0, if DistributionX is set to "GaussianZero" or "GaussianZeroCategorical".
24. `ShapeX.gamma`. This argument takes a numeric input, specifying the shape parameter k, if DistributionX is set to "Gamma", "GammaCategorical" or "NegativeBinomial". See also the rgamma() and rnbinom() functions for further details. Categorization is performed after the call to the rgamma() function.
25. `ScaleX.gamma`. This argument takes a numeric input, specifying the scale parameter capital theta, if DistributionX is set to "Gamma", "GammaCategorical" or "NegativeBinomial". See also the rgamma() and rnbinom() functions for further details. Categorization is performed after the call to the rgamma() function.
26. `DFX.student`. This argument takes a numeric input, specifying the degrees of freedom, if DistributionX is set to "StudentsT". See also the rt() function for further details.
27. `MinX.uni`. This argument takes a numeric input, specifying the minimum of the distribution, if DistributionX is set to "Uniform". See also the runif() function for further details.
28. `MaxX.uni`. This argument takes a numeric input, specifying the maximum of the distribution, if DistributionX is set to "Uniform". See also the runif() function for further details.
29. `LambdaX.pois`. This argument takes a numeric input, specifying the mean of the distribution, if DistributionX is set to "Poisson". See also the rpois() function for further details.

The function TrustGaussTypeII() can be used for the analysis of type II error rates as described in the main text. It adds a predefined effect to a single covariate only. All arguments can be accessed via

```
> ?TrustGaussTypeII
```

The function takes all the above 29 arguments of the TrustGauss() function and two additional ones.

30. `EffectXCov`. This argument takes a numeric input, specifying the effect size that should be simulated.
31. `ZTransform`. This argument is Boolean. If it is set to TRUE, the distributions of the dependent variable $Y$ and the independent variable $X$ are Z-transformed prior to adding the effect specified via EffectXCov.

The function TrustGaussLMM() can be used to fit linear mixed-effects models and to assess type I error rates. It takes the above 29 arguments of the TrustGauss() function. Furthermore, a single random effect can be specified via

32. `RanEF`. This argument is Boolean. If it is set to TRUE, a single random effect is fitted.
33. `nRanEFLevels`. This argument takes a numeric integer input, specifying how many repeated measures for each sampling points are generated.

34. `RanEFVarExp`. This argument takes a numeric input, specifying the amount of variance explained by the random effect. This amount is only correct if the variables are normally distributed.

The three functions in TrustGauss return a list with at least five elements

1. A data frame `Pvals` with all *P*-values of covariates and factors. It has as many rows as simulation runs (nSimulations) and as many columns as fitted covariates and factors. Each row represents the *P*-values of a single iteration. Column names are of the form "PCov1", "PCov2", …, "PFac1", "PFac2", … (depending on what combination of covariates and factors is specified). If CompareTtest was set to TRUE, a two column data frame is returned with columns "PFac1" and "PTtest". The second column represents the *P*-values of a *t*-test.
2. A data frame `ResCCC`. It has as many rows as simulation runs (nSimulations) and six columns. Each row represents the following estimates of a single iteration: Column "PSWY" contains all *P*-values of a Shapiro-Wilk test for normality of the dependent variable *Y*, column "PSWRes" contains all *P*-values of a Shapiro-Wilk test for normality of the residuals, column "rho" contains the concordance correlation coefficient (Lin 1989) between observed and expected residuals assuming normality for each model. We use the qqnorm() function to generate the expected values. Columns "s.shift", "l.shift" and "C.b" contain the scale shift, the location shift and the bias correction factor of the concordance correlation (Lin 1989). We use the CCC() function of the DescTools R package (v0.99.25; Signorell & mult. al. 2018) for estimating these parameters.
3. A vector `Alphas.05` that contains the type I error rate at an α-level of 0.05 for each of the fitted covariates and factors summarized across all simulation runs.
4. A vector `Alphas.001` that contains the type I error rate at an α-level of 0.001 for each of the fitted covariates and factors summarized across all simulation runs.
5. A data frame `ShapiroWilk`, which is a summary of the data frame ResCCC with one row and six columns. Column "Mean.PSWR" contains the mean *P*-value of the Shapiro-Wilk tests for normality of the residuals, calculated as

   $$10^{\texttt{mean(log10(ResCCC\$PSWRes))}}$$

   Columns "QL.PSWR" and "QU.PSWR" provide the lower and upper 95% quantiles of the *P*-values of the Shapiro-Wilk tests for normality of the residuals. Column "Mean.PSWY" contains the mean *P*-value of the Shapiro-Wilk tests for normality of the dependent variable *Y*, calculated as

   $$10^{\texttt{mean(log10(ResCCC\$PSWY))}}$$

   Columns "QL.PSWY" and "QU.PSWY" provide the lower and upper 95% quantiles of the *P*-values of the Shapiro-Wilk tests for normality of the dependent variable.

6. If the argument SaveAllOutput is set to TRUE, a list `Data` that contains as many data frames as there are dependent and independent variables fitted. The data frames are named "Dependent", "Cov1", "Cov2", … "Fac1", "Fac2", … (depending on what combination of covariates and factors is specified). In each of these data frames the values of *Y* (Dependent) and *X* (all the other data frames) are stored. Each of these data frames has as many rows as simulation runs (nSimulations) and as many columns as the specified sample size (nSamples). Each row represents the data values of a single iteration.

The function TrustGaussTypeII() returns one more element

7. A data frame `Effects` with the parameter estimates of the covariate with an added effect. It has as many rows as simulation runs (nSimulations) and three columns. Each row represents the parameter estimates of a single iteration. Column “Intercept” contains the estimate of the intercept, column “Estimate” contains the slope estimate and column “SE” the standard error of the slope estimate.

The function TrustGaussLMM() returns the same list as the TrustGauss() function with one additional element

8. A vector `VarExp` that contains the proportion of variance explained by the single random effect fitted in the mixed-effects model. Each element of the vector corresponds to a single iteration. Thus, it has as many elements as simulation runs (nSimulations).

**Description of the data generating functions**

All of the above three functions in the TrustGauss package generate data for the dependent variable *Y* and the independent variable *X* through calls to the base R functions rnorm(), rgamma(), rbinom(), rnbinom(), rt(), rpois(), runif() and sample(). These functions draw random values with specified parameter arguments from a Gaussian, Gamma, binomial, negative binomial, Student’s t, Poisson, uniform (floating numbers) or uniform (integers) distribution, respectively. The distributions “GaussianCategorical”, “GaussianZero”, “GaussianZeroCategorical”, “AbsoluteGaussian” and “GammaCategorical” make use of additional functions to generate categories, to take absolute values or to introduce zero-inflation after a call to rnorm() or rgamma(), thereby changing the specified mean and standard deviation or shape and scale. Table 1 lists the specific parameter settings for the ten distributions simulated in the main text (D0–D9). Figures 1A and S2A display histograms of the distributions D0–D9.

**A typical call to TrustGauss()**

Assume we want to run a simulation with 100 observations in each of 50,000 iterations. We specify the Family argument as “Gaussian” to fit a linear model with identity link. See the glm() documentation for details on the Family arguments. This is equivalent to fitting a linear model assuming normally distributed errors. Since we might also be interested in the individual observations of each simulation run, we specify SaveAllOutput=TRUE. This will save the 100 × 50,000 = 5,000,000 data points of the dependent variable *Y* and the 5,000,000 observations of the predictor *X*. We want the dependent variable *Y* to be distributed as the absolute values of a Gaussian distribution with mean 0 and standard deviation 1. The independent variable *X* should be a single covariate that is Gamma distributed with shape 1.5 and scale 10. Thus, our call to TrustGauss() looks like this

```
> sim <- TrustGauss(Family="gaussian", nSamples=100,
  nSimulations=50000, SaveAllOutput=TRUE,
  DistributionY="AbsoluteGaussian", DistributionXCov="Gamma",
  DistributionXFac=NULL, MeanY.gauss=0, SDY.gauss=1,
  ShapeX.gamma=1.5, ScaleX.gamma=10)
```

A progress bar is going to indicate the progress of the simulation, which is updated after every iteration. As soon as the simulation has finished, we can access every element of the resulting list. For example, if we want to obtain the data frame of individual *P*-values for every iteration in this call to TrustGauss(), we can access it via

```
> sim$Pval
```

A summary of the type I error rate at an α-level of 0.05 is accessible through

```
> sim$Alphas.05
```

In this call to TrustGauss() with a single covariate, this will yield only a single value for the independent variable *X*. If we had multiple covariates or factors specified, a separate type I error rate for each of them would have been displayed.

**Description of the basic simulations**

After specifying the input arguments for the TrustGauss()function (see above), the simulation starts by generating uncorrelated data for the dependent variable *Y* and the independent variable *X* according to the input parameters. Following our above example, specifying a sample size of nSamples = 100 and the distribution of *Y* as DistributionY = "AbsoluteGaussian" with MeanY.gauss = 0 and SDY.gauss = 1, the function generates data for *Y* as

```
> y <- abs(rnorm(n=100, mean=0, sd=1))
```

Similarly, for the independent variable *X* we specify DistributionX = "Gamma" with ShapeX.gamma = 1.5 and ScaleX.gamma = 10. Then, the function generates data for *X* as

```
> x <- rgamma(n=100, shape=1.5, scale=10)
```

After the data generating step, two linear models are fitted using the Family argument (argument 1) from above

```
> mod1 <- glm(y ~ x, family=Family)
> mod2 <- glm(y ~ 1, family=Family)
```

The models are compared via

```
> anova(mod1, mod2)
```

We keep the *P*-value of this model comparison and start the next iteration by generating data for *Y* and *X* again. The number of iterations was set to nSimulations = 50,000 in all our simulations. We further set Family = "gaussian", with the exception of models with a Poisson error structure that are highlighted as such in the main text.

Because the data for *Y* and *X* are uncorrelated, we expect 5% of all models to yield a *P*-value ≤ 0.05. If more than 5% of all models have a *P*-value ≤ 0.05, then the type I error rate is inflated (i.e. too many models yield "significant" results), whereas if less models have a *P*-value ≤ 0.05, then they are conservative, yielding too few "significant" results. In Figure 1, combinations of *Y* and *X* that produce inflated type I error rates are coloured red and those yielding conservative *P*-values are coloured blue.

**Introduction of heteroscedasticity**

First, we sampled the independent variable *X* from a binomial distribution, where we varied the success rate from 0.2 to 0.8 in steps of 0.1. Whenever the independent variable *X* was equal to 0, we sampled the dependent variable *Y* either from distribution D0 or D7 (see Table 1). We also introduced a third distribution D7.1, which was negative binomial with mean 0.5 and a variance of 1 (with the rational of introducing the same absolute difference in variance as in D0). Whenever the independent variable *X* was equal to 1, we sampled the dependent variable from the same distribution but with a 10-times larger variance. We assessed the effects of heteroscedasticity with sample sizes of $N = 100$ and $N = 1000$. We then fitted a glm either with a Gaussian or a Quasipoisson error structure, where we tested the significance of the independent variable *X* via a likelihood ratio test. We fitted these models to 50,000 datasets for each combination of the dependent and predictor variable with two sample sizes (i.e. 3 distribution of the dependent variable × 7 distributions of the independent variable × 2 sample sizes = 42 combinations, each with a Gaussian or a Quasipoisson error structure).

Second, we introduced a second independent variable *X* that we sampled from a uniform distribution with five levels. The other predictor and the dependent variable *Y* were sampled as described above with sample sizes of $N = 100$ and $N = 1000$. We then fitted a glm either with a Gaussian or a Quasipoisson error structure, where we tested the significance of the interaction between the two independent variables via a likelihood ratio test. We fitted these models to 50,000 datasets for each combination of the dependent and predictor variables with two sample sizes as well.

For each simulation run, we recorded the variance in the two groups (as defined by the predictor variable with two levels, see above). We summarized the 50,000 simulation runs by calculating (1) the type I error rate as the number of simulations with a $P$-value ≤ 0.05 / the number of all simulation runs (i.e. 50,000) and (2) the mean observed difference in variances between the two groups (see Table S5).

### Supplementary Figures

**Figure S1 |** All simulated combinations of the dependent variable *Y* and the predictor *X* that were fitted in linear regression models for sample sizes *N* = 10, 25, 50, 100, 250, 500, 1000. Figure names are constructed as “X” <Distribution name of *X*> “_Y” <How many distributions of *Y*> “Distributions_N” <Sample size> “_Sim” <Number of simulation runs>, such that the file “XD0_Y10Distributions_N10_Sim50000” shows results from all 50,000 simulation runs where the predictor *X* was normally distributed, the independent variable *Y* had ten different distributions (D0–D9) and the sample size was *N* = 10. In each of the 7 × 10 = 70 figures, the leftmost column depicts the distribution of *Y*, the second column depicts the distribution of *X*, the third column a QQ-plot of the residuals and the forth column a QQ-plot of the -$\log_{10}$(*P*-values). Residuals were distributed as the dependent variable *Y* because the regression coefficient *b* was on average zero.

*Provided as supplementary figures (.jpg) on the Open Science Framework homepage.*

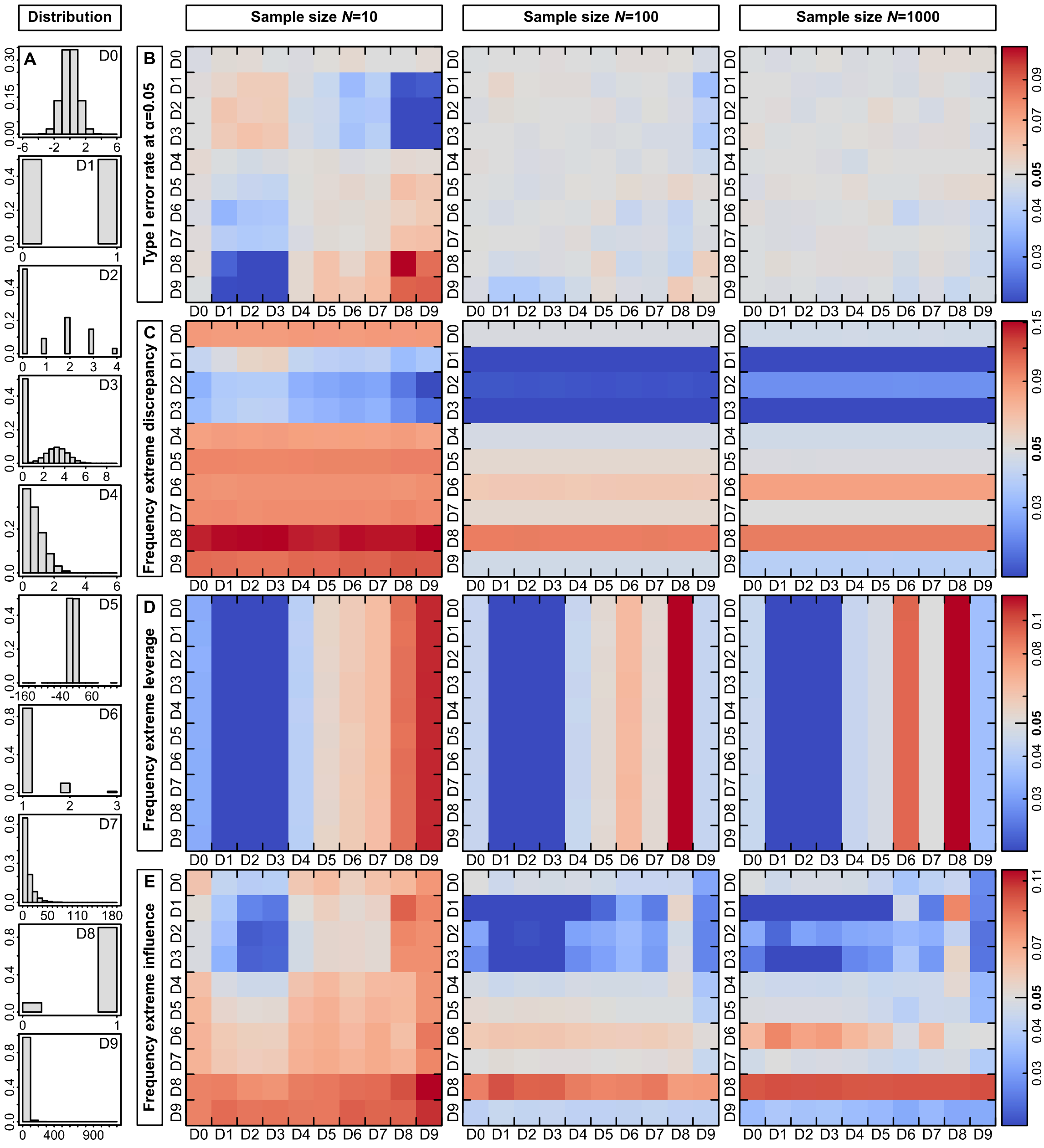


**Figure S2 |** Summary of linear model diagnostics for all 10 × 10 = 100 combinations of the dependent variable *Y* and the predictor *X* depicted in (**A**). The numbers D0–D9 refer to the plots in in (**B–E**) where on the *Y*-axis the distribution of the dependent variable and on the *X*-axis of the predictor is indicated. (**B**) Type I error rate at an α-level of 0.05 for sample sizes of $N = 10$, 100 and 1000. Red colours represent increased and blue conservative type I error rates. (**C**) Mean proportion of studentized residuals (*R*) exceeding the critical value of $R > 2$ as a measure of discrepancy. A large discrepancy value represents an observation whose dependent variable *Y* is unusual given its value of the predictor *X*. It is thus influenced predominately by the distribution of *Y*. (**D**) Mean proportion of hat values (*H*) exceeding the critical value of $H > (2 \times (k + 1)) / n$ as a measure of leverage. *k* is the number of regression slopes and *n* is the number of observations. A large leverage value represents an observation whose predictor *X* is unusual given its value of the independent variable *Y*. It is thus influenced predominately by the distribution of *X*. (**E**) Mean proportion of Cook's distance (*D*) exceeding the critical value of $D > 4 / (n - k - 1)$ as a measure of influence. Influence represents the product of discrepancy and leverage (Zuur, Ieno & Smith 2007; Ramsey & Schafer 2013).

**Figure S3 |** All simulated combinations of the dependent variable *Y* and the last of four predictors *X* that were fitted in linear regression models. The first three predictors were normally distributed and the distribution of the last one was varied. The sample size was *N* = 100. For a description of file names and content see Figure S1.

*Provided as supplementary figures (.jpg) on the Open Science Framework homepage.*

**Figure S4 |** All simulated combinations of the dependent variable *Y* and the predictor *X* fitted in linear random-intercept models. We simulated $N$ = 100 independent samples each of which was sampled twice, such that the single random effect explained roughly 30% of the variation in *Y*. For a description of file names and content see Figure S1.

*Provided as supplementary figures (.jpg) on the Open Science Framework homepage.*

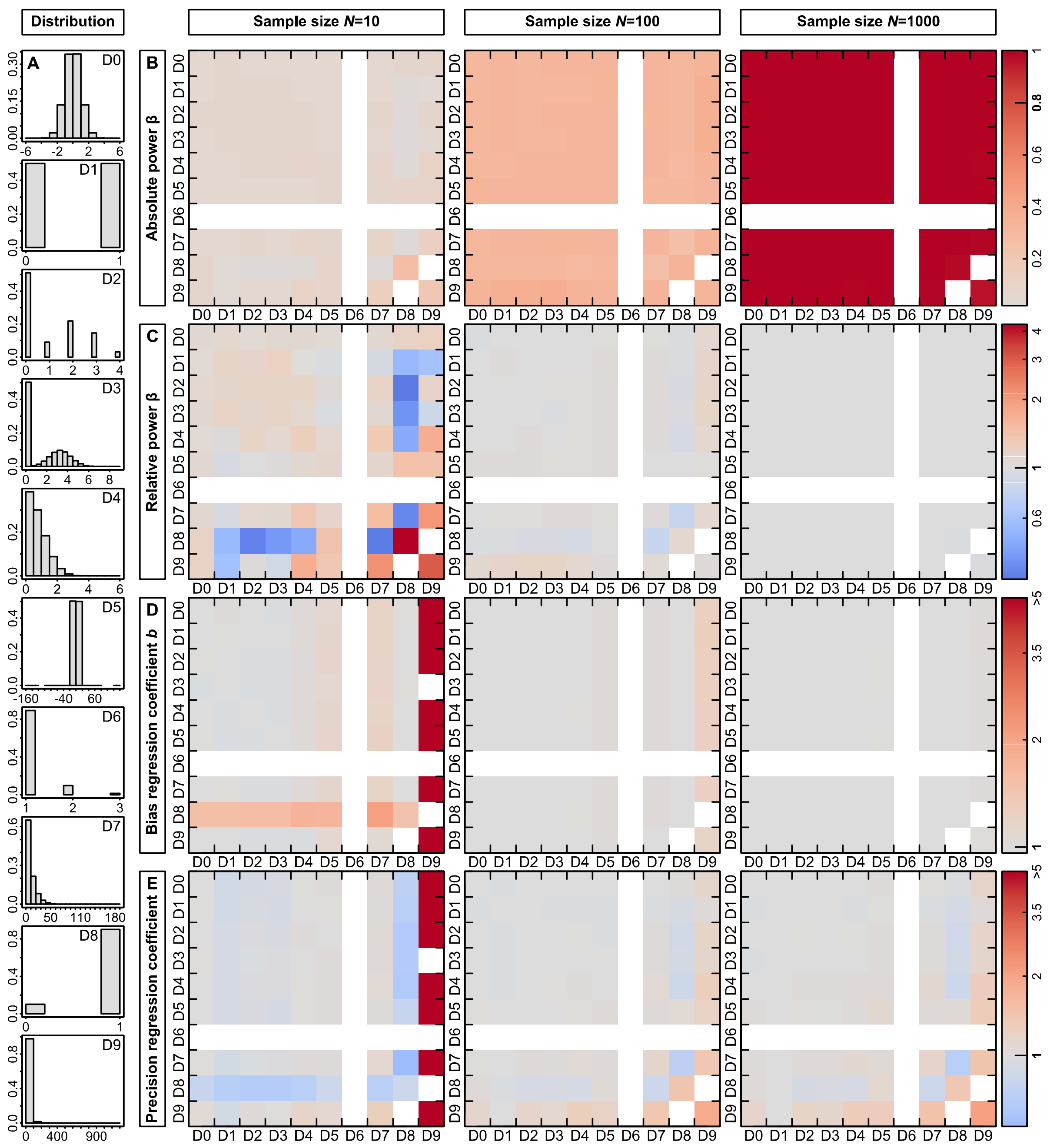


**Figure S5 |** Power, bias and precision of parameter estimates from Gaussian linear regression models for all 10 × 10 = 100 combinations of the dependent variable *Y* and the predictor *X* at a regression coefficient *b* = 0.15. (**A**) Overview of the different distributions that we simulated, which were the same as in Figure 1. The numbers D0–D9 refer to the plots in (**B**–**E**) where on the *Y*-axis the distribution of the dependent variable and on the *X*-axis of the predictor is indicated. (**B**) Power for sample sizes of *N* = 10, 100 and 1000. Red colours represent increased power. (**C**) Deviation of power from the expected value derived from a normally distributed *Y* and *X* for sample sizes of *N* = 10, 100 and 1000. Red colours represent increased and blue colours decreased power. (**D**) Bias and (**E**) precision of the regression coefficient estimates for sample sizes of *N* = 10, 100 and 1000.

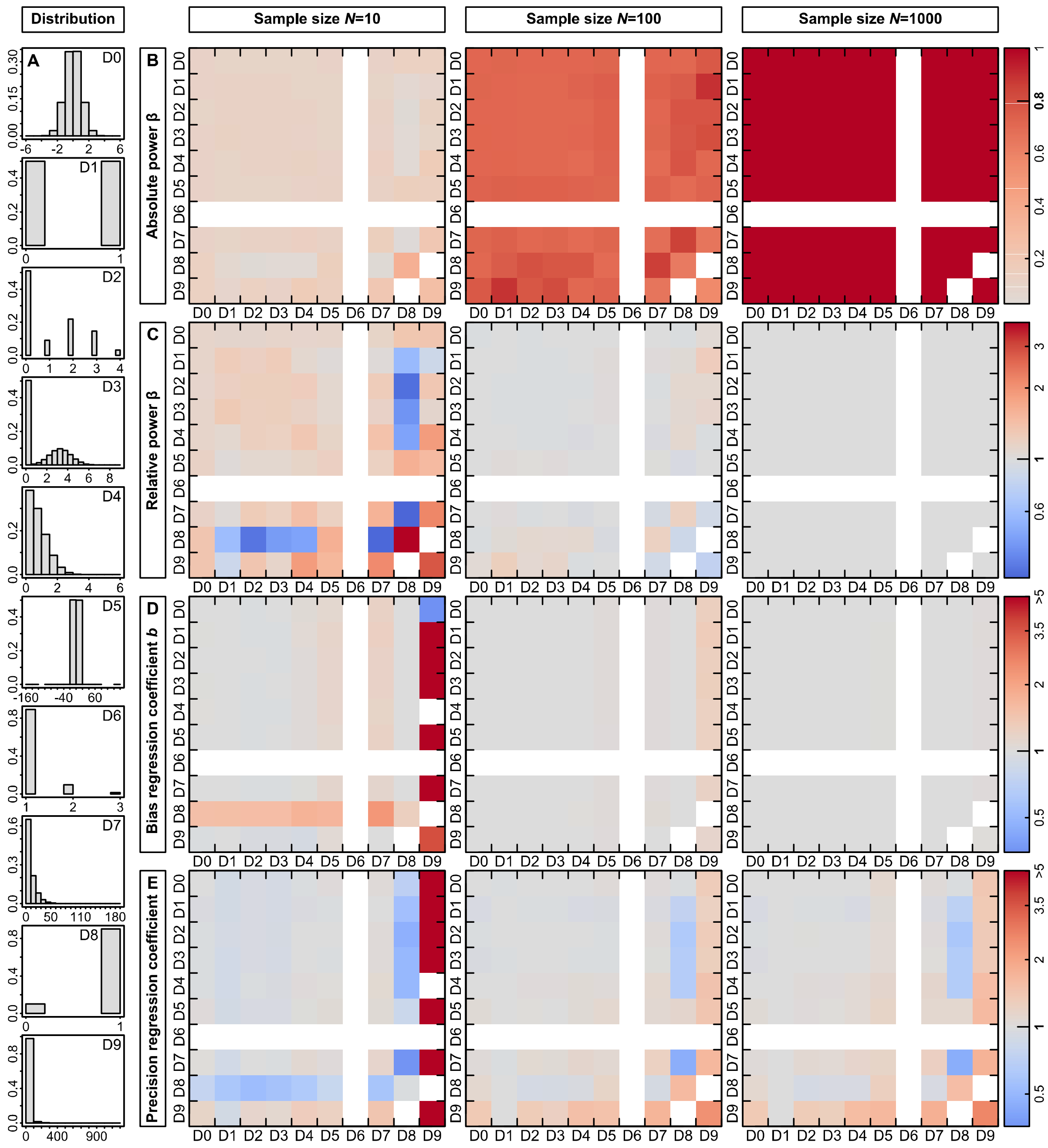


**Figure S6 |** Power, bias and precision of parameter estimates from Gaussian linear regression models for all 10 × 10 = 100 combinations of the dependent variable *Y* and the predictor *X* at a regression coefficient *b* = 0.25. (**A**) Overview of the different distributions that we simulated, which were the same as in Figure 1. The numbers D0–D9 refer to the plots in (**B**–**E**) where on the *Y*-axis the distribution of the dependent variable and on the *X*-axis of the predictor is indicated. (**B**) Power for sample sizes of *N* = 10, 100 and 1000. Red colours represent increased power. (**C**) Deviation of power from the expected value derived from a normally distributed *Y* and *X* for sample sizes of *N* = 10, 100 and 1000. Red colours represent increased and blue colours decreased power. (**D**) Bias and (**E**) precision of the regression coefficient estimates for sample sizes of *N* = 10, 100 and 1000.

**Figure S7 |** All simulated combinations of the dependent variable *Y* and the predictor *X* fitted in generalized linear models with a Poisson error structure. The sample size was $N = 100$. For a description of file names and content see Figure S1.

*Provided as supplementary figures (.jpg) on the Open Science Framework homepage.*

### Supplementary Tables

**Table S1 |** Observed and expected power of a regression model in which both the dependent variable *Y* and the predictor *X* are normally distributed with mean 0 and standard deviation 1. The expected power was calculated using the power.SLR() function from the powerMediation R package (v0.2.9, Dupont & Plummer 1998; Qiu 2018). The observed power was estimated using 50,000 simulations.

| Sample size | Mean of slope *b* | Expected power at $\alpha = 0.05$ | Expected power at $\alpha = 0.001$ | Observed power at $\alpha = 0.05$ | Observed power at $\alpha = 0.001$ |
|---:|---:|---:|---:|---:|---:|
| 10 | 0.15 | 0.064 | $1.20 \times 10^{-3}$ | 0.069 | $1.48 \times 10^{-3}$ |
| 10 | 0.20 | 0.077 | $1.38 \times 10^{-3}$ | 0.084 | $2.26 \times 10^{-3}$ |
| 10 | 0.25 | 0.094 | $1.64 \times 10^{-3}$ | 0.106 | $3.44 \times 10^{-3}$ |
| 100 | 0.15 | 0.321 | 0.032 | 0.316 | 0.034 |
| 100 | 0.20 | 0.523 | 0.090 | 0.517 | 0.095 |
| 100 | 0.25 | 0.724 | 0.210 | 0.714 | 0.218 |
| 1000 | 0.15 | 0.998 | 0.933 | 0.998 | 0.932 |
| 1000 | 0.20 | 1.000 | 0.999 | 1.000 | 0.999 |
| 1000 | 0.25 | 1.000 | 1.000 | 1.000 | 1.000 |

**Table S2 |** Distributions and effect sizes used for assessing the interpretability and power of Gaussian versus binomial and Poisson models at a sample size of *N* = 100 in 50,000 simulation runs. Each combination of the dependent variable *Y* and predictor *X* was fitted in a glm using a Gaussian error structure and the appropriate error structure according to the distribution of *Y*. The effect sizes were chosen such that we reached a power of around 0.5.

| Sampling distribution *Y* | Sampling distribution *X* | Mean *Y* | Variance *Y* | Mean *X* | Variance *X* | Effect | Back-transformed effect[#] |
|---|---|---|---|---|---|---|---|
| Poisson | Gaussian | 1 | 1 | 0 | 1 | 0.2 | 1.22 |
| Poisson | Gamma | 1 | 1 | 10 | 100 | 2.2 | 9.03 |
| Poisson | Binomial | 1 | 1 | 0.75 | 0.19 | 0.1 | 1.11 |
| Binomial | Gaussian | 0.75 | 0.19 | 0 | 1 | 0.45 | 0.61 |
| Binomial | Gamma | 0.75 | 0.19 | 10 | 100 | 4.2 | 0.99 |
| Binomial | Binomial | 0.75 | 0.19 | 0.75 | 0.19 | 0.2 | 0.55 |

### Back-transformation was done by using the functions plogis() for binomial models and exp() for Poisson models.

**Table S3 |** Summary of type I error rates (α), scale shift parameters (υ), mean observed *P*-values at an expected value of $10^{-3}$ and $10^{-4}$ and the resulting bias from 50,000 simulation runs across all 10 × 10 = 100 combinations of the dependent variable *Y* and the predictor *X*.

| **Sample size** | **Mean α (range)** | **Mean υ (range)** | **Mean *P* at $10^{-3}$ (range)** | **Mean *P* at $10^{-4}$ (range)** | **Bias at $-\log_{10}P=3$** | **Bias at $-\log_{10}P=4$** |
|---|---|---|---|---|---|---|
| 10 | 0.048 (0.015–0.110) | 2.437 (0.696–51.350) | $7.72 \times 10^{-12}$ ($2.59 \times 10^{-125}$–$7.00 \times 10^{-3}$) | $7.96 \times 10^{-21}$ ($6.31 \times 10^{-129}$–$1.05 \times 10^{-3}$) | 3.704 (0.718–41.529) | 5.025 (0.744–32.050) |
| 25 | 0.048 (0.019–0.073) | 1.065 (0.771–2.461) | $1.96 \times 10^{-4}$ ($2.20 \times 10^{-16}$–$7.93 \times 10^{-3}$) | $3.25 \times 10^{-6}$ ($6.81 \times 10^{-27}$–$1.57 \times 10^{-3}$) | 1.236 (0.700–5.219) | 1.372 (0.701–6.542) |
| 50 | 0.048 (0.028–0.067) | 1.043 (0.837–2.229) | $3.64 \times 10^{-4}$ ($5.36 \times 10^{-13}$–$6.47 \times 10^{-3}$) | $5.94 \times 10^{-6}$ ($3.55 \times 10^{-23}$–$1.27 \times 10^{-3}$) | 1.146 (0.730–4.090) | 1.307 (0.724–5.612) |
| 100 | 0.049 (0.037–0.058) | 1.026 (0.892–1.937) | $5.04 \times 10^{-4}$ ($8.22 \times 10^{-11}$–$4.55 \times 10^{-3}$) | $1.76 \times 10^{-5}$ ($7.14 \times 10^{-19}$–$1.45 \times 10^{-3}$) | 1.099 (0.781–3.362) | 1.188 (0.710–4.537) |
| 250 | 0.049 (0.041–0.053) | 1.020 (0.941–1.631) | $5.81 \times 10^{-4}$ ($2.49 \times 10^{-9}$–$3.00 \times 10^{-3}$) | $2.16 \times 10^{-5}$ ($2.14 \times 10^{-15}$–$3.31 \times 10^{-4}$) | 1.079 (0.841–2.868) | 1.166 (0.870–3.667) |
| 500 | 0.049 (0.045–0.052) | 1.014 (0.958–1.534) | $6.57 \times 10^{-4}$ ($9.12 \times 10^{-9}$–$2.02 \times 10^{-3}$) | $3.12 \times 10^{-5}$ ($1.87 \times 10^{-16}$–$7.07 \times 10^{-4}$) | 1.061 (0.898–2.680) | 1.126 (0.788–3.932) |
| 1000 | 0.049 (0.044–0.052) | 1.010 (0.977–1.343) | $7.51 \times 10^{-4}$ ($6.35 \times 10^{-7}$–$1.52 \times 10^{-3}$) | $3.94 \times 10^{-5}$ ($1.29 \times 10^{-14}$–$2.52 \times 10^{-4}$) | 1.042 (0.939–2.066) | 1.101 (0.899–3.473) |

**Table S4 |** Summary of type I error rates (α), scale shift parameters (υ), mean observed *P*-values at an expected value of $10^{-3}$ and $10^{-4}$ and the resulting bias from 50,000 simulation runs across all 10 × 2 = 20 combinations where either the dependent variable *Y* or the predictor *X* was normally distributed.

| Normally distributed variable | Sample size | Mean α (range) | Mean υ (range) | Mean *P* at $10^{-3}$ (range) | Mean *P* at $10^{-4}$ (range) | Bias at $-\log_{10}P$=3 | Bias at $-\log_{10}P$=4 |
|---|---|---|---|---|---|---|---|
| Y | 10 | 0.050 (0.048–0.052) | 1.005 (0.997–1.015) | $8.82 \times 10^{-4}$ ($6.64 \times 10^{-4}$–$1.03 \times 10^{-3}$) | $8.43 \times 10^{-5}$ ($4.01 \times 10^{-5}$–$1.55 \times 10^{-4}$) | 1.018 (0.996–1.059) | 1.019 (0.952–1.099) |
| Y | 25 | 0.050 (0.049–0.052) | 1.004 (0.990–1.015) | $9.90 \times 10^{-4}$ ($7.92 \times 10^{-4}$–$1.22 \times 10^{-3}$) | $6.25 \times 10^{-5}$ ($3.19 \times 10^{-5}$–$1.17 \times 10^{-4}$) | 1.002 (0.971–1.034) | 1.051 (0.983–1.124) |
| Y | 50 | 0.050 (0.049–0.051) | 0.999 (0.992–1.015) | $1.02 \times 10^{-3}$ ($8.06 \times 10^{-4}$–$1.32 \times 10^{-3}$) | $9.14 \times 10^{-5}$ ($4.79 \times 10^{-5}$–$1.55 \times 10^{-4}$) | 0.997 (0.960–1.031) | 1.010 (0.952–1.080) |
| Y | 100 | 0.050 (0.048–0.051) | 1.000 (0.993–1.005) | $1.02 \times 10^{-3}$ ($8.90 \times 10^{-4}$–$1.25 \times 10^{-3}$) | $8.22 \times 10^{-5}$ ($5.90 \times 10^{-5}$–$1.32 \times 10^{-4}$) | 0.998 (0.968–1.017) | 1.021 (0.970–1.057) |
| Y | 250 | 0.050 (0.048–0.051) | 1.001 (0.990–1.014) | $9.66 \times 10^{-4}$ ($7.49 \times 10^{-4}$–$1.22 \times 10^{-3}$) | $8.29 \times 10^{-5}$ ($3.77 \times 10^{-5}$–$1.61 \times 10^{-4}$) | 1.005 (0.972–1.042) | 1.020 (0.949–1.106) |
| Y | 500 | 0.049 (0.047–0.051) | 0.998 (0.984–1.003) | $9.95 \times 10^{-4}$ ($7.62 \times 10^{-4}$–$1.28 \times 10^{-3}$) | $8.39 \times 10^{-5}$ ($4.20 \times 10^{-5}$–$1.79 \times 10^{-4}$) | 1.001 (0.965–1.039) | 1.019 (0.937–1.094) |
| Y | 1000 | 0.050 (0.049–0.052) | 0.999 (0.989–1.017) | $1.06 \times 10^{-3}$ ($8.99 \times 10^{-4}$–$1.36 \times 10^{-3}$) | $8.47 \times 10^{-5}$ ($5.86 \times 10^{-5}$–$1.44 \times 10^{-4}$) | 0.992 (0.956–1.015) | 1.018 (0.960–1.058) |
| X | 10 | 0.050 (0.048–0.052) | 1.003 (0.996–1.014) | $9.28 \times 10^{-4}$ ($7.59 \times 10^{-4}$–$1.24 \times 10^{-3}$) | $9.76 \times 10^{-5}$ ($5.62 \times 10^{-5}$–$1.46 \times 10^{-4}$) | 1.011 (0.968–1.040) | 1.003 (0.959–1.063) |
| X | 25 | 0.050 (0.049–0.051) | 0.999 (0.988–1.010) | $1.11 \times 10^{-3}$ ($8.45 \times 10^{-4}$–$1.42 \times 10^{-3}$) | $9.73 \times 10^{-5}$ ($5.31 \times 10^{-5}$–$1.69 \times 10^{-4}$) | 0.986 (0.949–1.024) | 1.003 (0.943–1.069) |
| X | 50 | 0.049 (0.048–0.052) | 0.998 (0.988–1.017) | $1.01 \times 10^{-3}$ ($7.53 \times 10^{-4}$–$1.27 \times 10^{-3}$) | $1.35 \times 10^{-4}$ ($7.49 \times 10^{-5}$–$2.03 \times 10^{-4}$) | 0.998 (0.965–1.041) | 0.968 (0.923–1.031) |
| X | 100 | 0.050 (0.048–0.051) | 1.001 (0.996–1.005) | $9.98 \times 10^{-4}$ ($8.04 \times 10^{-4}$–$1.20 \times 10^{-3}$) | $7.64 \times 10^{-5}$ ($2.39 \times 10^{-5}$–$1.81 \times 10^{-4}$) | 1.000 (0.973–1.032) | 1.029 (0.936–1.155) |
| X | 250 | 0.050 (0.049–0.052) | 1.004 (0.995–1.016) | $9.69 \times 10^{-4}$ ($7.49 \times 10^{-4}$–$1.25 \times 10^{-3}$) | $6.74 \times 10^{-5}$ ($2.72 \times 10^{-5}$–$1.46 \times 10^{-4}$) | 1.005 (0.968–1.042) | 1.043 (0.959–1.141) |
| X | 500 | 0.050 (0.049–0.052) | 1.000 (0.989–1.014) | $9.67 \times 10^{-4}$ ($7.99 \times 10^{-4}$–$1.16 \times 10^{-3}$) | $1.15 \times 10^{-4}$ ($5.19 \times 10^{-5}$–$1.84 \times 10^{-4}$) | 1.005 (0.979–1.033) | 0.985 (0.934–1.071) |
| X | 1000 | 0.050 (0.049–0.052) | 1.000 (0.992–1.012) | $1.02 \times 10^{-3}$ ($7.48 \times 10^{-4}$–$1.30 \times 10^{-3}$) | $6.92 \times 10^{-5}$ ($1.91 \times 10^{-5}$–$1.38 \times 10^{-4}$) | 0.997 (0.962–1.042) | 1.040 (0.965–1.180) |

**Table S5 |** Summary of the heteroscedasticity simulations. We estimated type I error rates (α) in glms with a Gaussian or Quasipoisson error structure and the mean observed difference in variances between the two groups as defined by the predictor variable with two levels (see Supplementary Methods for details, expected value is 10).

| Model | Sampling distribution $Y$ | P($X_1$=1) | Sample size | Gaussian α | Quasipoisson α | Observed difference in variances |
|---|---|---|---|---|---|---|
| $Y \sim X_1$ | D0 | 0.2 | 100 | $1.18 \times 10^{-3}$ | - | 11.21 |
| $Y \sim X_1$ | D0 | 0.3 | 100 | $7.70 \times 10^{-3}$ | - | 10.75 |
| $Y \sim X_1$ | D0 | 0.4 | 100 | 0.024 | - | 10.53 |
| $Y \sim X_1$ | D0 | 0.5 | 100 | 0.054 | - | 10.41 |
| $Y \sim X_1$ | D0 | 0.6 | 100 | 0.101 | - | 10.35 |
| $Y \sim X_1$ | D0 | 0.7 | 100 | 0.168 | - | 10.31 |
| $Y \sim X_1$ | D0 | 0.8 | 100 | 0.262 | - | 10.24 |
| $Y \sim X_1$ | D7 | 0.2 | 100 | 0.037 | 0.024 | 14.46 |
| $Y \sim X_1$ | D7 | 0.3 | 100 | 0.063 | 0.047 | 12.78 |
| $Y \sim X_1$ | D7 | 0.4 | 100 | 0.096 | 0.080 | 11.97 |
| $Y \sim X_1$ | D7 | 0.5 | 100 | 0.130 | 0.121 | 11.73 |
| $Y \sim X_1$ | D7 | 0.6 | 100 | 0.176 | 0.180 | 11.37 |
| $Y \sim X_1$ | D7 | 0.7 | 100 | 0.237 | 0.256 | 11.17 |
| $Y \sim X_1$ | D7 | 0.8 | 100 | 0.319 | 0.356 | 10.95 |
| $Y \sim X_1$ | D7.1 | 0.2 | 100 | 0.110 | 0.079 | 19.21 |
| $Y \sim X_1$ | D7.1 | 0.3 | 100 | 0.145 | 0.119 | 15.86 |
| $Y \sim X_1$ | D7.1 | 0.4 | 100 | 0.177 | 0.158 | 14.39 |
| $Y \sim X_1$ | D7.1 | 0.5 | 100 | 0.201 | 0.199 | 13.66 |
| $Y \sim X_1$ | D7.1 | 0.6 | 100 | 0.219 | 0.244 | 13.12 |
| $Y \sim X_1$ | D7.1 | 0.7 | 100 | 0.207 | 0.283 | 13.42 |
| $Y \sim X_1$ | D7.1 | 0.8 | 100 | 0.154 | 0.298 | 15.00 |
| $Y \sim X_1$ | D0 | 0.2 | 1000 | $5.4 \times 10^{-4}$ | - | 10.10 |
| $Y \sim X_1$ | D0 | 0.3 | 1000 | $6.68 \times 10^{-3}$ | - | 10.07 |
| $Y \sim X_1$ | D0 | 0.4 | 1000 | 0.022 | - | 10.05 |
| $Y \sim X_1$ | D0 | 0.5 | 1000 | 0.050 | - | 10.04 |
| $Y \sim X_1$ | D0 | 0.6 | 1000 | 0.097 | - | 10.03 |
| $Y \sim X_1$ | D0 | 0.7 | 1000 | 0.166 | - | 10.03 |
| $Y \sim X_1$ | D0 | 0.8 | 1000 | 0.250 | - | 10.02 |
| $Y \sim X_1$ | D7 | 0.2 | 1000 | $4.34 \times 10^{-3}$ | $2.80 \times 10^{-3}$ | 10.39 |
| $Y \sim X_1$ | D7 | 0.3 | 1000 | 0.015 | 0.012 | 10.25 |
| $Y \sim X_1$ | D7 | 0.4 | 1000 | 0.031 | 0.028 | 10.20 |
| $Y \sim X_1$ | D7 | 0.5 | 1000 | 0.062 | 0.059 | 10.15 |

| | | | | | | |
|---|---|---|---|---|---|---|
| $Y \sim X_1$ | D7 | 0.6 | 1000 | 0.109 | 0.108 | 10.13 |
| $Y \sim X_1$ | D7 | 0.7 | 1000 | 0.174 | 0.176 | 10.09 |
| $Y \sim X_1$ | D7 | 0.8 | 1000 | 0.265 | 0.270 | 10.11 |
| $Y \sim X_1$ | D7.1 | 0.2 | 1000 | 0.015 | $8.56 \times 10^{-3}$ | 10.74 |
| $Y \sim X_1$ | D7.1 | 0.3 | 1000 | 0.030 | 0.021 | 10.49 |
| $Y \sim X_1$ | D7.1 | 0.4 | 1000 | 0.056 | 0.046 | 10.38 |
| $Y \sim X_1$ | D7.1 | 0.5 | 1000 | 0.089 | 0.081 | 10.24 |
| $Y \sim X_1$ | D7.1 | 0.6 | 1000 | 0.132 | 0.131 | 10.19 |
| $Y \sim X_1$ | D7.1 | 0.7 | 1000 | 0.197 | 0.203 | 10.17 |
| $Y \sim X_1$ | D7.1 | 0.8 | 1000 | 0.288 | 0.299 | 10.21 |
| $Y \sim X_1 * X_2$ | D0 | 0.2 | 100 | $1.2 \times 10^{-4}$ | - | 11.23 |
| $Y \sim X_1 * X_2$ | D0 | 0.3 | 100 | $2.42 \times 10^{-3}$ | - | 10.76 |
| $Y \sim X_1 * X_2$ | D0 | 0.4 | 100 | 0.017 | - | 10.58 |
| $Y \sim X_1 * X_2$ | D0 | 0.5 | 100 | 0.064 | - | 10.47 |
| $Y \sim X_1 * X_2$ | D0 | 0.6 | 100 | 0.169 | - | 10.34 |
| $Y \sim X_1 * X_2$ | D0 | 0.7 | 100 | 0.340 | - | 10.29 |
| $Y \sim X_1 * X_2$ | D0 | 0.8 | 100 | 0.559 | - | 10.27 |
| $Y \sim X_1 * X_2$ | D7 | 0.2 | 100 | $4.26 \times 10^{-3}$ | $9.20 \times 10^{-3}$ | 14.52 |
| $Y \sim X_1 * X_2$ | D7 | 0.3 | 100 | $7.40 \times 10^{-3}$ | 0.036 | 12.83 |
| $Y \sim X_1 * X_2$ | D7 | 0.4 | 100 | 0.019 | 0.097 | 12.09 |
| $Y \sim X_1 * X_2$ | D7 | 0.5 | 100 | 0.047 | 0.206 | 11.58 |
| $Y \sim X_1 * X_2$ | D7 | 0.6 | 100 | 0.108 | 0.353 | 11.26 |
| $Y \sim X_1 * X_2$ | D7 | 0.7 | 100 | 0.209 | 0.492 | 11.07 |
| $Y \sim X_1 * X_2$ | D7 | 0.8 | 100 | 0.321 | 0.557 | 11.09 |
| $Y \sim X_1 * X_2$ | D7.1 | 0.2 | 100 | 0.049 | 0.084 | 20.10 |
| $Y \sim X_1 * X_2$ | D7.1 | 0.3 | 100 | 0.042 | 0.147 | 15.97 |
| $Y \sim X_1 * X_2$ | D7.1 | 0.4 | 100 | 0.036 | 0.217 | 14.06 |
| $Y \sim X_1 * X_2$ | D7.1 | 0.5 | 100 | 0.044 | 0.293 | 13.57 |
| $Y \sim X_1 * X_2$ | D7.1 | 0.6 | 100 | 0.080 | 0.357 | 13.20 |
| $Y \sim X_1 * X_2$ | D7.1 | 0.7 | 100 | 0.150 | 0.381 | 13.50 |
| $Y \sim X_1 * X_2$ | D7.1 | 0.8 | 100 | 0.251 | 0.368 | 15.17 |
| $Y \sim X_1 * X_2$ | D0 | 0.2 | 1000 | $2 \times 10^{-5}$ | - | 10.11 |
| $Y \sim X_1 * X_2$ | D0 | 0.3 | 1000 | $1.00 \times 10^{-3}$ | - | 10.07 |
| $Y \sim X_1 * X_2$ | D0 | 0.4 | 1000 | 0.011 | - | 10.05 |
| $Y \sim X_1 * X_2$ | D0 | 0.5 | 1000 | 0.051 | - | 10.04 |
| $Y \sim X_1 * X_2$ | D0 | 0.6 | 1000 | 0.146 | - | 10.03 |
| $Y \sim X_1 * X_2$ | D0 | 0.7 | 1000 | 0.306 | - | 10.02 |
| $Y \sim X_1 * X_2$ | D0 | 0.8 | 1000 | 0.526 | - | 10.04 |

| | | | | | | |
|---|---|---|---|---|---|---|
| $Y \sim X_1 * X_2$ | D7 | 0.2 | 1000 | $4 \times 10^{-5}$ | $1.4 \times 10^{-4}$ | 10.42 |
| $Y \sim X_1 * X_2$ | D7 | 0.3 | 1000 | $1.24 \times 10^{-3}$ | $3.24 \times 10^{-3}$ | 10.25 |
| $Y \sim X_1 * X_2$ | D7 | 0.4 | 1000 | $9.82 \times 10^{-3}$ | 0.020 | 10.21 |
| $Y \sim X_1 * X_2$ | D7 | 0.5 | 1000 | 0.046 | 0.077 | 10.16 |
| $Y \sim X_1 * X_2$ | D7 | 0.6 | 1000 | 0.136 | 0.195 | 10.13 |
| $Y \sim X_1 * X_2$ | D7 | 0.7 | 1000 | 0.297 | 0.379 | 10.09 |
| $Y \sim X_1 * X_2$ | D7 | 0.8 | 1000 | 0.509 | 0.596 | 10.06 |
| $Y \sim X_1 * X_2$ | D7.1 | 0.2 | 1000 | $1.4 \times 10^{-4}$ | $1.12 \times 10^{-3}$ | 10.71 |
| $Y \sim X_1 * X_2$ | D7.1 | 0.3 | 1000 | $1.26 \times 10^{-3}$ | 0.010 | 10.49 |
| $Y \sim X_1 * X_2$ | D7.1 | 0.4 | 1000 | $8.78 \times 10^{-3}$ | 0.046 | 10.34 |
| $Y \sim X_1 * X_2$ | D7.1 | 0.5 | 1000 | 0.037 | 0.128 | 10.34 |
| $Y \sim X_1 * X_2$ | D7.1 | 0.6 | 1000 | 0.115 | 0.278 | 10.24 |
| $Y \sim X_1 * X_2$ | D7.1 | 0.7 | 1000 | 0.259 | 0.482 | 10.25 |
| $Y \sim X_1 * X_2$ | D7.1 | 0.8 | 1000 | 0.449 | 0.684 | 10.19 |
